# An Anaerobic Pathogen Rewires Host Metabolism to Fuel Oxidative Growth in the Inflamed Gut

**DOI:** 10.1101/2025.05.31.657111

**Authors:** Luisella Spiga, Ryan T. Fansler, Alexandra Grote, Madison Langford-Butler, Asia K. Miller, Maxwell Neal, Owen F. Hale, Yifan Wu, Deepanshu Singla, M. Wade Calcutt, Abigail E. Rose, Madeline M. Bresson, Alexandra C. Schrimpe-Rutledge, Brittany Berdy, Simona G. Codreanu, Mary Kay Washington, Benjamin P. Bratton, Stacy D. Sherrod, John A. McLean, Karsten Zengler, Cynthia L. Sears, Megan G. Behringer, Andreas Gnirke, Jonathan Livny, Danyvid Olivares-Villagómez, Ashlee M. Earl, Wenhan Zhu

## Abstract

To colonize their host and cause disease, enteric pathogens must deploy their virulence factors to establish distinct nutrient niches. How obligate anaerobic pathogens construct nutrient niches in the densely populated large intestine remains poorly understood. Enterotoxigenic *Bacteroides fragilis* (ETBF) is considered an obligate anaerobic bacterium and has been implicated in inflammation-associated diseases, including colitis and colorectal cancer. Here we show that ETBF uses its virulence factor, *Bacteroides fragilis* toxin, to reprogram colonic epithelial cell metabolism to colonize the inflamed gut. *Bacteroides fragilis* toxin activates colonic epithelial signaling and hijacks the host bile acid recycling pathway, inducing a metabolic shift in the epithelium from oxidative phosphorylation to glycolysis. This shift increases local concentrations of lactate and oxygen, nutrients that support an oxidative metabolism in ETBF. These findings reveal an unexpected strategy by which a pathogenic organism, previously considered to be an obligate anaerobic bacterium, generates and exploits an oxidative niche in the inflamed gut.

## Introduction

Enteric infections are a leading cause of acute diarrheal diseases and have recently been implicated in the development of chronic conditions such as colorectal cancer (CRC)^1, 2^. CRC develops at the largest host-microbe interface in the human body^3^ and is predominantly sporadic, with most cases lacking well-defined host germline mutations^4, 5^. Consequently, environmental factors, particularly exposure to enteric pathogens, are increasingly recognized as key contributors to CRC pathogenesis^6–10^. Despite these associations, the mechanisms by which enteric pathogens colonize the gut and influence disease progression remain poorly understood.

Enterotoxigenic *Bacteroides fragilis* (ETBF), a toxin-producing subclass of the common gut commensal *B. fragilis*, has been implicated in both inflammatory diarrheal and CRC across independent human studies^11–24^. ETBF also promotes inflammatory colitis^25^ and colonic tumorigenesis in gnotobiotic^26^ or conventional^16, 27–30^ murine models. These pathogenic effects are primarily driven by the virulence factor *Bacteroides fragilis* toxin (BFT), which elicits a range of physiological alterations in host cells^27, 31–36^. However, the specific mechanisms by which BFT facilitates ETBF niche establishment and promotes persistent colonization in the gut remain largely undefined.

Colonocyte metabolism is central to host-microbe interactions as it shapes the composition and functions of the gut microbiome^37^. In the healthy gut, colonocytes generate energy through β-oxidation of microbiota-derived short-chain fatty acids, such as butyrate^38, 39^. This process consumes oxygen from the underlying mucosal surface, effectively maintaining an anaerobic environment in the colonic lumen^40^. As a result, obligate anaerobic bacteria, which rely on anaerobic fermentation to generate energy, dominate the gut microbiota^41^. To successfully colonize the densely populated intestine, pathogens must employ virulence factors to manipulate host responses, carving out new nutrient niches^42^. For instance, facultative anaerobes such as *Salmonella enterica serovar* Typhimurium and *Citrobacter rodentium* subvert colonocyte metabolism to increase luminal oxygen availability^43–45^. This strategy enables them to engage in respiratory metabolism^43–45^, thereby outcompeting the resident anaerobic bacteria that rely on the energetically less-efficient anaerobic fermentation to grow^37^.

In contrast to these facultative anaerobic pathogens, members of the *Bacteroidetes* phylum, including ETBF (*B. fragilis*), are classically viewed as obligate anaerobes that are either killed or cease growing upon oxygen exposure^46, 47^. As such, they reside in anoxic niches, such as the mammalian colon, and primarily generate energy via anaerobic fermentation of dietary or host-derived glycans^48, 49^. Current dogma holds that these obligate anaerobes, including ETBF, lack α-ketoglutarate dehydrogenase (SucAB), succinyl-CoA synthetase (SucCD), and succinate dehydrogenase (Sdh). This results in a bifurcation of the TCA cycle^50^, incomplete oxidation of acetyl-CoA, and, consequently, less efficient energy production than a fully oxidative TCA cycle. How ETBF, long considered an anaerobic pathogen, successfully persists in the inflamed, oxygen-enrich gut it induces^51^, has remained unclear. Here, we demonstrate that ETBF leverages its virulence factor BFT to reprogram epithelial cell metabolism, thereby reshaping the gut nutritional landscape. This reprogramming leads to increased levels of lactate and oxygen, which fuel ETBF’s unique oxidative metabolism. These findings challenge the long-standing view of *B. fragilis* as strictly anaerobic and uncover a novel mechanism by which an obligate anaerobe reshapes its environment to establish an oxidative metabolic niche and promote its own expansion in the inflamed gut.

## Results

### Anaerobe ETBF operates an oxidative metabolism to respire oxygen

Since most nutritional niches in the gut are already occupied by the resident microbiota, enteric pathogens often employ virulence strategies to elicit host response, facilitating the establishment of novel niches that enable their colonization^43–45, 52–56^. To explore whether BFT, a recognized virulence factor previously shown to induce a series of inflammatory changes, reshapes the intestinal nutritional landscape to facilitate ETBF colonization, we colonized groups of C57BL/6 mice with the ETBF wild-type strain and the isogenic toxin-deficient Δ*bft* mutant and assessed their fitness and host inflammatory status. Indeed, loss of *bft* markedly dampened inflammation and reduced pathogen load compared to the wild-type counterpart (**Fig. 1A-C**).

**Fig. 1.**
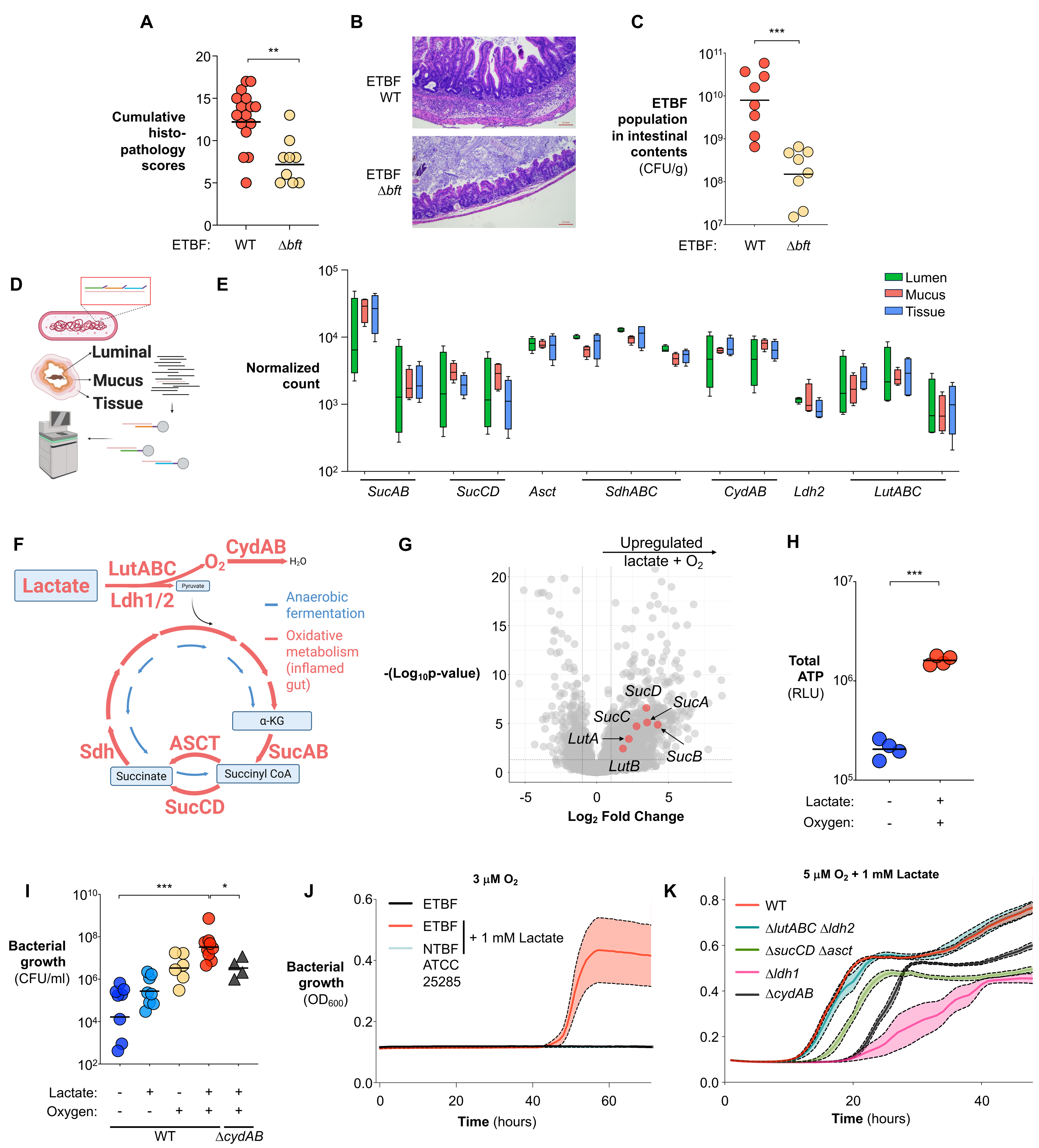
ETBF engages oxidative metabolism during intestinal colonization. (**A-E**) Groups of C57BL/6 mice were colonized with either wild-type ETBF or an isogenic Δ*bft* mutant for 7 days. (**A**) Histopathology scores of ETBF-infected animals. (**B**) Representative H&E-stained images of intestinal tissue. (**C**) ETBF populations in intestinal contents were quantified by plating on selective media. Hybrid-selection RNA-seq was performed on luminal contents, mucus, and cecal tissue to profile the ETBF transcriptome during infection. Transcriptomic data were integrated into a genome-scale metabolic model. (**D**) Schematic overview of the Hybrid-selection RNA-seq approach. (**E**) Normalized transcript counts of indicated metabolic genes across sample sites. (**F**) Predicted metabolic flux through genes involved in lactate oxidation, oxygen respiration, and oxidative central metabolism during ETBF infection, based on transcript abundance (highlighted in red). (**G**) Wild-type ETBF was subcultured in BHI medium supplemented with 1 mM lactate under microaerobic conditions (1 μM O_2_). Differentially expressed genes relative to control conditions were visualized by volcano plot. (**H-K**) Indicated ETBF strains were cultured in media supplemented with 1 mM lactate and exposed to the indicated oxygen levels. (**H**) Total cellular ATP levels. (**I-K**) *In vitro* growth of ETBF strains under the indicated conditions. In panels **A, C, E, H** and **I**, bars represent geometric means. In panels **J** and **K**, connected lines and shaded areas represent the mean and SEM. *, *p*<0.05; ***, *p*<0.001.

Next, we sought to determine how ETBF adapts to the inflammatory niches sculpted by BFT. Because bacteria tailor their transcriptional programs to the available nutrients, the bacterial transcriptome can serve as a proxy for the functional state of ETBF metabolism *in vivo*^57, 58^. However, pathogen mRNA is vastly outnumbered by host transcripts^59^ and those from the diverse commensal bacteria^60, 61^. To overcome this, we employed hybrid selection RNA-sequencing (hsRNA-seq)^62^, using capture probes tiled across all predicted ETBF open-reading frames. This approach enables the selective capture of ETBF transcripts from total RNA isolated from host tissues and gut microbial community without introducing substantial enrichment bias (**Fig. 1D and S1A**). Transcriptomic analysis by RNA-Seq without target capture revealed that host tissues mount robust responses to BFT, likely reshaping the gut nutritional landscape (**Fig. S1C**). In turn, ETBF adapts its transcriptome to the nutritional landscape altered by BFT, with 35 pathways significantly enriched among the differentially expressed genes in the lumen (**Fig. S1D**). Notably, these pathways include “ribosome”, “gene expression”, and “translation”, all of which are energy-demanding processes, suggesting that BFT-induced host responses create a nutrient niche that enhances ETBF fitness in the inflamed gut.

Strikingly, analysis of the genomic contents revealed that ETBF encodes key enzymes (Sdh, SucAB, SucCD) necessary for a complete TCA cycle (**Fig. 1E**). ETBF also codes for acetate: succinate CoA transferase (Asct)^63^, which catalyzes succinyl-CoA to succinate conversion similar to SucCD, but with the added production of acetyl-CoA (**Fig. 1F**), further supporting ETBF’s capacity to operate an oxidative central metabolism. Because these genes are dispensable for fermentative growth and are generally suppressed under anaerobic conditions^64, 65^, their expression could reflect adaptation to oxygen availability. Remarkably, we observed robust transcription of *sdh*, *sucAB*, *sucCD*, and *asct* in ETBF-colonized murine intestines (**Fig. 1E&F**), challenging the long-held view that *B. fragilis* is an obligate anaerobe. With notable exceptions of *sdh* and *asct*, transcript levels of oxidative metabolic genes were generally highest in bacteria associated with intestinal tissue, compared to those in mucus or lumen fractions (**Fig. 1E**), a possible reflection of the oxygen gradient that decreases toward the lumen. Integrating the *in vivo* transcriptome into genome-scale metabolic modeling predicted substantial flux through three lactate dehydrogenases (Ldh1, Ldh2 (RS13195), LutABC), along with components of oxygen respiration (CydAB) and oxidative TCA cycle (SucAB, SucCD, Asct) (**Fig. 1F**). In contrast, the fluxes through these genes are minimal under anaerobic conditions (**Fig. 1F**). Interestingly, *sucAB* (E1 subunit), *sucCD*, and *ldh2* display mosaic distribution within the *Bacteroides* genus, with higher prevalence in those members with relatively higher oxygen-tolerance, suggesting these genes may participate in adaptation to oxidative niches^66^ (**Fig. S1E**). Together, these findings suggest that ETBF operates a metabolic program that diverges from the strictly anaerobic fermentation pathways typically associated with other *Bacteroides*^67^.

### Oxygen and lactate drive an oxidative metabolism in ETBF

Cellular respiration involves the oxidation of an electron donor (e.g., lactate) and the transfer of the resulting electrons through an electron transport chain to an electron acceptor, such as oxygen^68^. By allowing for more complete oxidation of carbon sources and more efficient energy conservation, respiration generates ATP more efficiently, conferring a metabolic advantage over anaerobic fermentation^69^. Oxygen is the most favorable electron acceptor, given the high free energy yield; however, most obligate anaerobes are killed by oxygen or derived reactive oxygen species (ROS), precluding its use for respiration^46, 70^. Notably, recent work revealed that a subset of anaerobes (nanaerobe), including *B. fragilis*, exhibit enhanced oxygen tolerance^71^ and can use nanomolar concentrations of oxygen as an electron acceptor for additional fitness boosts^66, 72, 73^. However, the mechanisms that enable oxygen respiration in nanaerobes and their physiological relevance within the mammalian host remain largely unknown. In facultative anaerobes, the expression levels of the TCA cycle enzymes primarily respond to the presence of oxygen and carbon sources, such as lactate^64, 74, 75^. The abundant transcripts of genes involved in lactate utilization suggest that lactate may be a candidate carbon source for ETBF in the inflamed gut (**Fig. 1E**). We thus asked whether oxygen and lactate drive a complete TCA cycle in ETBF. *In vitro* exposure to oxygen and lactate upregulated genes involved in lactate oxidation and oxidative central metabolism (**Fig. 1G**). As an excellent electron acceptor, oxygen enables the utilization of otherwise poorly fermentable carbon sources, such as lactate^76^, thereby facilitating ATP production^77^. Consistent with this, oxygen and lactate synergistically increased both ATP production and bacterial fitness of ETBF *in vitro* (**Fig. 1H-I**).

To further test this hypothesis, we inoculated ETBF at low density (2.5x10^4^ CFU) into an open, micro-anaerobic atmosphere with 3-10 μM O_2_ (0.3-1%) to minimize the impact of collective scavenging^78^ of oxygen that can occur at high bacterial loads and the resulting decrease in oxygen tension due to microbial consumption in sealed vessles^66^. Remarkably, lactate supported ETBF growth in the presence of up to 10 μM oxygen, an order of magnitude higher concentration than the nanomolar concentrations previously reported for nanaerobes^66, 72, 73^, in a rich, semi-defined medium where other carbon sources are available (**Fig. 1J&S1F**). At 5 µM O_2_, ETBF entered exponential phase earlier than under anaerobic conditions, consistent with enhanced metabolic efficiency (**Fig. S1F**). In contrast, a non-toxigenic *B. fragilis* strain (ATCC 25285) failed to benefit from these nutrients even at 3 µM O_2_ (**Fig. 1J**). Notably, the key enzymes that support oxidative metabolism in ETBF are largely conserved in the non-toxigenic strain, suggesting that differences in transcriptional regulation or enzyme function may underlie the distinct metabolic adaptations to oxygenated environments between the two strains.

Next, we tested whether ETBF carries out oxidative central metabolism that could enable the coupling of electron-donating lactate oxidation to electron-consuming oxygen respiration^76, 79, 80^. Indeed, deleting the genes required for oxygen respiration (11*cydAB*)^46^, lactate utilization (Δ*ldh1* or 11*lutABC* 11*ldh2*; note that repeated attempts to generate the triple mutant were unsuccessful), or oxidative central metabolism (11*sucCD* 11*asct*) in ETBF significantly impaired its ability to benefit from lactate oxidation under micro-aerobic conditions (**Fig. 1K**), but had no influence on bacterial fitness when ETBF was cultured anaerobically (**Fig. S2A**). Together, this data suggests that ETBF, despite being classed as an “anaerobe”, operates an oxidative metabolism to couple lactate oxidation and oxygen respiration.

### ETBF operates an oxidative metabolism to colonize the intestine

BFT plays an important role in facilitating ETBF colonization (**Fig. 1A-C**), likely by establishing a niche optimized for the pathogen’s fitness. This observation raised the question of how ETBF exploits BFT-mediated changes for gut colonization. Given that ETBF couples lactate oxidation to oxygen respiration *in vitro*, and actively transcribes genes involved in oxidative metabolism in the inflamed gut (**Fig. 1**), we hypothesize that ETBF relies on oxidative metabolism to exploit the environment created by BFT for efficient gut colonization. Consistent with this notion, ablating the terminal oxidase^46^ (11*cydAB*) in the oxygen respiration pathway significantly attenuated ETBF colonization and partially reduced infection-associated intestinal inflammation (**Fig. 2A-D**). Weighted Gene Co-expression Network Analysis (WGCNA) of the intestinal tissue transcriptome revealed that the Δ*cydAB* mutant exhibited reduced infection-associated changes in host processes such as epithelial-mesenchymal transition, reinforcing that ETBF alters host processes to reshape gut nutritional environment (**Fig. S2B-C**).

**Fig. 2.**
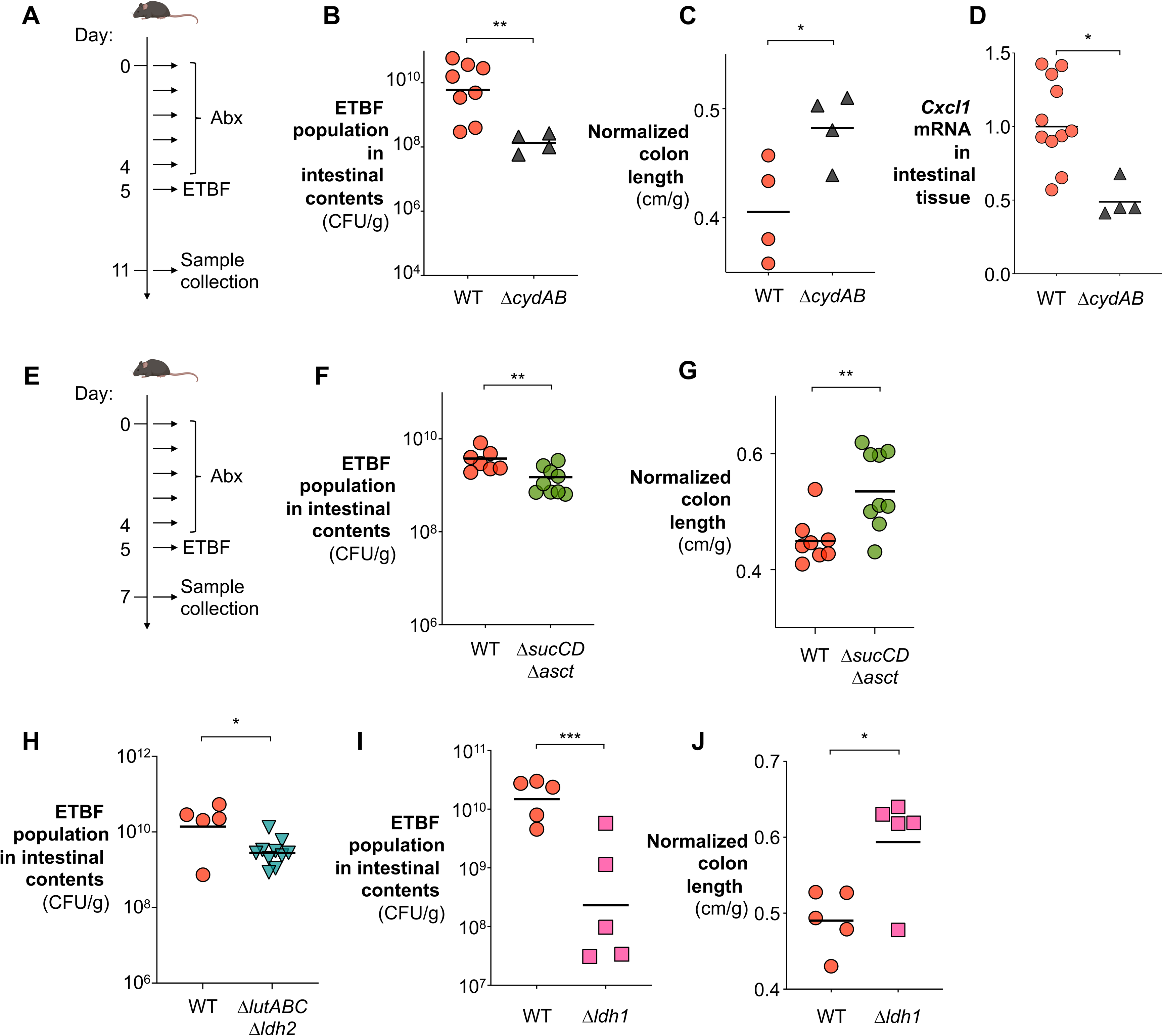
Oxygen respiration and oxidative metabolism contribute to ETBF colonization and inflammation. (**A-D**) Groups of C57BL/6 mice were colonized with the indicated ETBF strains for 7 days. (**A**) Schematic representation of the experiment. (**B**) ETBF populations in intestinal contents quantified by plating on selective media. (**C**) Colon length normalized to body weight. (**D**) Cxcl1 transcript levels in intestinal tissue measured by RT-qPCR. (**E-J**) C57BL/6 mice were colonized with the indicated ETBF strains for 3 days. (**E**) Schematic representation of the experiment. (**F, H, I**) ETBF populations in intestinal contents quantified by plating on selective media. (**G, J**) Colon length normalized to body weight. Bars represent geometric means. *, *p*<0.05; **, *p*<0.01; ***, *p*<0.001.

Engaging an oxidative central metabolism could enable ETBF to couple lactate oxidation with oxygen respiration through an electron transport chain^76, 81^, facilitating its gut colonization. To test this hypothesis, we assessed the *in vivo* fitness of mutants lacking lactate dehydrogenases (Ldh1, LutABC & Ldh2) and key oxidative TCA cycle components (SucCD & Asct, SucAB). In agreement with our *in vitro* observations (**Fig. 1K**), these genes were critical for ETBF fitness and pathogenicity *in vivo* (**Fig. 2E-J**). In particular, the Δ*ldh1* mutant exhibited an even more pronounced defect in colonizing the intestinal mucus layer, suggesting that lactate utilization in this oxygenated niche is a pathogen-adaptive strategy (**Fig. S2D**). In contrast, deletion of *sucAB* did not result in a notable fitness cost, indicating possible functional redundancy provided by an alternative, uncharacterized enzyme (**Fig. S2E**).

Notably, the reduced colonization observed in the Δ*sucCD* Δ*asct* and Δ*ldh1* mutants was accompanied by attenuated intestinal inflammation (**Fig. 2G, J; Fig. S2F**), whereas deletion of the other two lactate dehydrogenases (Δ*lutABC* Δ*ldh2*) or *sucAB* alone had no impact on the inflammatory response compared to the wild-type strain (**Fig. S2G&H**). Together, these findings demonstrate that oxygen respiration, oxidative metabolism, and selective lactate oxidation are key metabolic drivers of ETBF fitness in the inflamed gut, revealing a previously unappreciated capacity of this obligate anaerobe to exploit oxygenated inflammatory niches.

### BFT rewires colonocyte metabolism, increasing oxygen and lactate levels in the gut

Previous studies have shown that BFT elicits a range of host epithelial changes that may converge to reshape the intestinal nutritional landscape. These include the induction of cleavage of zonula adherens junction protein E-cadherin^82^ and hyperplasia through dysregulating Wnt/β-catenin/c-Myc signaling^25, 83^. Crypt hyperplasia, in turn, expands the population of immature colonocytes, which preferentially engage in anaerobic glycolysis rather than the oxidative phosphorylation typical of differentiated colonocytes^84, 85^. Beyond direct effects on epithelial cells, BFT also triggers pathological activation of T-helper 17 cells (T_H_17)^27, 31^, which could further exacerbate the switch to glycolysis through the effector cytokine IL-17^86, 87^. Collectively, these effects drive a shift from oxidative metabolism to glycolysis in colonocytes^37, 44, 85, 88–92^, thereby increasing lactate export and allowing oxygen to diffuse from the lamina propria into the lumen^80, 93^. Consistent with this hypothesis, RNA-seq of colonic tissue of ETBF-infected mice revealed BFT-dependent repression of genes involved in β-oxidation and electron-transport-chain complexes I-IV (β-oxidation, electron transport chain complex I-IV) (**Fig. 3A**).

**Fig. 3.**
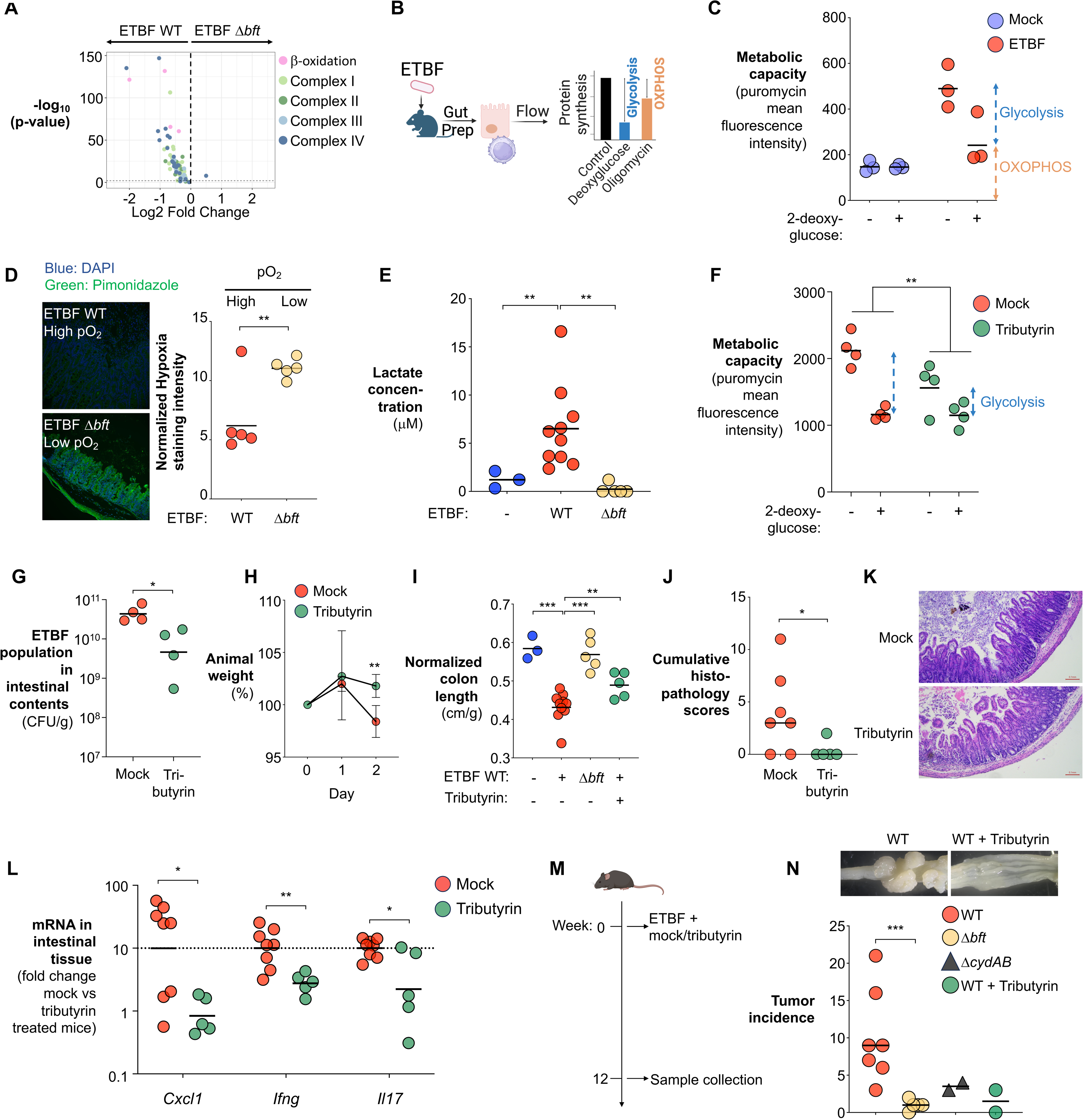
BFT-driven metabolic reprogramming of colonocytes promotes ETBF colonization. (**A-E**) Groups of C57BL/6 mice were colonized with the indicated ETBF strains for 7 days. (**A**) RNA-seq analysis of intestinal tissues revealed BFT-dependent repression of genes involved in colonocyte oxidative metabolism (β-oxidation, electron transport chain Complexes I–IV). (**B-C**) Metabolic capacity and glycolytic dependency of gut epithelial cells were profiled using the SCENITH assay. (**B**) Schematic of the SCENITH method. (**C**) Quantification of metabolic activity and glycolytic dependency in gut epithelial cells. (**D**) Mice were injected intraperitoneally with pimonidazole 1 hour before euthanasia. Hypoxia was assessed by immunofluorescence using a-pimonidazole antiserum and Alexa-488-conjugated secondary antibody (green fluorescence), with DAPI counterstaining of nuclei (blue fluorescence). Hypoxia staining intensity was normalized to nuclear signal. (**E**) Lactate concentrations in intestinal contents were measured by LC-MS. (**F-L**) ETBF-colonized C57BL/6 mice were treated with tributyrin or vehicle control for 7 days. (**F**) Quantification of metabolic activity and glycolytic dependency in gut epithelial cells using SCENITH assay. (**G**) ETBF populations in intestinal contents were quantified by plating on selective media. (**H**) Animal weight. (**I**) Colon length normalized to body weight. (**J**) Cumulative histopathology scores based on H&E-stained intestinal tissue. (**K**) Representative H&E-stained images of intestinal tissue. (**L**) Transcript levels of inflammatory markers were determined by RT-qPCR. (**M-N**) *Apc*^Min^ mice were colonized with the indicated ETBF strains and received a single dose of tributyrin or a mock control. Samples were collected 12 weeks post-inoculation. (**M**) Experimental schematic. (**N**) Total tumor counts in the colon. Bars represent geometric means. **, *p*<0.01; ***, *p*<0.001.

To further assess the metabolic state of colonocytes during ETBF infection, we employed SCENITH assay to profile metabolic responses in various cell types *ex vivo* using flow cytometry^94^. SCENITH leverages puromycin incorporation into nascent proteins to measure protein synthesis levels, which serve as a reliable indicator of global metabolic activity^95^. By using inhibitors of oxidative phosphorylation and glycolysis, we could evaluate the relative contributions of these pathways (**Fig. 3B**). In agreement with previous reports^84, 85, 93^ and our transcriptomic analysis (**Fig. 3A**), colonocytes primarily operate an oxidative metabolism at homeostasis (**Fig. 3C**). However, upon ETBF infection, colonocytes underwent a notable metabolic shift toward glycolysis (**Fig. 3C**).

A shift from oxidative to glycolytic metabolism in colonocytes reduces oxygen consumption from the underlying vasculature, allowing oxygen to diffuse into the gut lumen^93, 96^. Concomitantly, lactate is secreted as a metabolic end product^97, 98^. Elevated luminal oxygen levels can also impair oxygen-sensitive enzymes in obligate anaerobic commensals, disrupting key metabolic pathways, including central carbon metabolism^99^. This oxidative stress may force commensals to reroute pyruvate toward lactate production^100^, further contributing to lactate accumulation in the gut. Indeed, measurement of intestinal oxygenation using pimonidazole, a compound that forms irreversible adducts with proteins or DNA in hypoxic tissues^101^, revealed that ETBF infection led to increased oxygen tension (**Fig. 3D**). Additionally, lactate quantification revealed a parallel rise in luminal lactate in a BFT-dependent manner, suggesting the shift to anaerobic glycolysis in colonocytes creates an oxidative, lactate-rich niche (**Fig. 3E**).

If heightened oxygen and lactate foster ETBF growth, restoring epithelial oxidative metabolism should counteract infection. To test this, we therapeutically administered tributyrin, a short-chain fatty acid precursor that promotes oxidative metabolism in colonocytes^93^, two days post-ETBF inoculation. Feature-level clustering (**Fig. S3A**), Weighted Gene Co-expression Network Analysis (WGCNA) analysis (**Fig. S3B**), and gene set enrichment analysis (**Fig. S3C**) revealed that tributyrin suppressed glycolytic gene expression. Consistent with these transcriptional changes, tributyrin partially reversed the ETBF-induced metabolic reprogramming of colonocytes toward glycolysis (**Fig. 3F**) and significantly reduced ETBF colonization (**Fig. 3G**). Furthermore, tributyrin treatment ameliorated ETBF-induced weight loss, attenuated intestinal inflammation, and mitigated associated histopathological changes (**Fig. 3H-L**). Of note, tributyrin did not exhibit direct antibacterial activity against ETBF (**Fig. S3D**) or significantly alter microbiota composition (**Fig. S3E-I**), indicating that its protective effect derives from modulation of host metabolism rather than microbial interference. Collectively, these findings support a model in which ETBF exploits elevated luminal oxygen to establish and sustain colonization during gut inflammation.

We next examined whether operating an oxidative metabolism facilitates ETBF-induced tumorigenesis. For this purpose, we utilized the multiple intestinal neoplasia (Min) (heterozygous for the adenomatous polyposis coli (*Apc*) gene) mouse model. Although *Apc*^Min^ mice are considered a small intestine tumorigenesis model, independent studies have demonstrated that ETBF colonization converts *Apc*^Min^ mice into a colon tumor model^27, 102–104^. Notably, ETBF-infected *Apc*^Min^ mice develop colonic tumors within weeks, well before small intestinal adenomas are clinically evident or detectable^105^. To disrupt ETBF’s respiratory metabolism, we either: (1) genetically ablated the terminal oxidase complex (Δ*cydAB*), thereby disrupting ETBF’s ability to respire, and (2) pharmacologically restored epithelial hypoxia by administering a single dose of tributyrin (**Fig. 3M**). Twelve weeks post-infection, we assessed colonic tumor incidence. Consistent with our hypothesis, both strategies significantly reduced tumor burden (**Fig. 3N**). These findings support a model in which ETBF-induced tumorigenesis is potentiated by its capacity to exploit oxygen and engage in oxidative metabolism, suggesting a link between microbial respiration and cancer promotion in the gut.

### BFT rewires colonocyte metabolism through interference with epithelial processes

A hallmark feature of BFT activity is the induction of proteolytic cleavage of the tight junction protein E-cadherin^34, 82^, compromising the colonic barrier integrity. Consistently, administration of FITC-dextran revealed ETBF markedly increased gut permeability (**Fig. 4A**), which could potentially allow oxygen diffusion across the gut epithelium to fuel ETBF colonization^37^. Supporting this, treatment with Marimastat^106^, an inhibitor of matrix metalloproteases capable of cleaving E-cadherin^107^, reduced ETBF colonization and ameliorated animal weight loss (**Fig. 4B-C**).

**Fig. 4.**
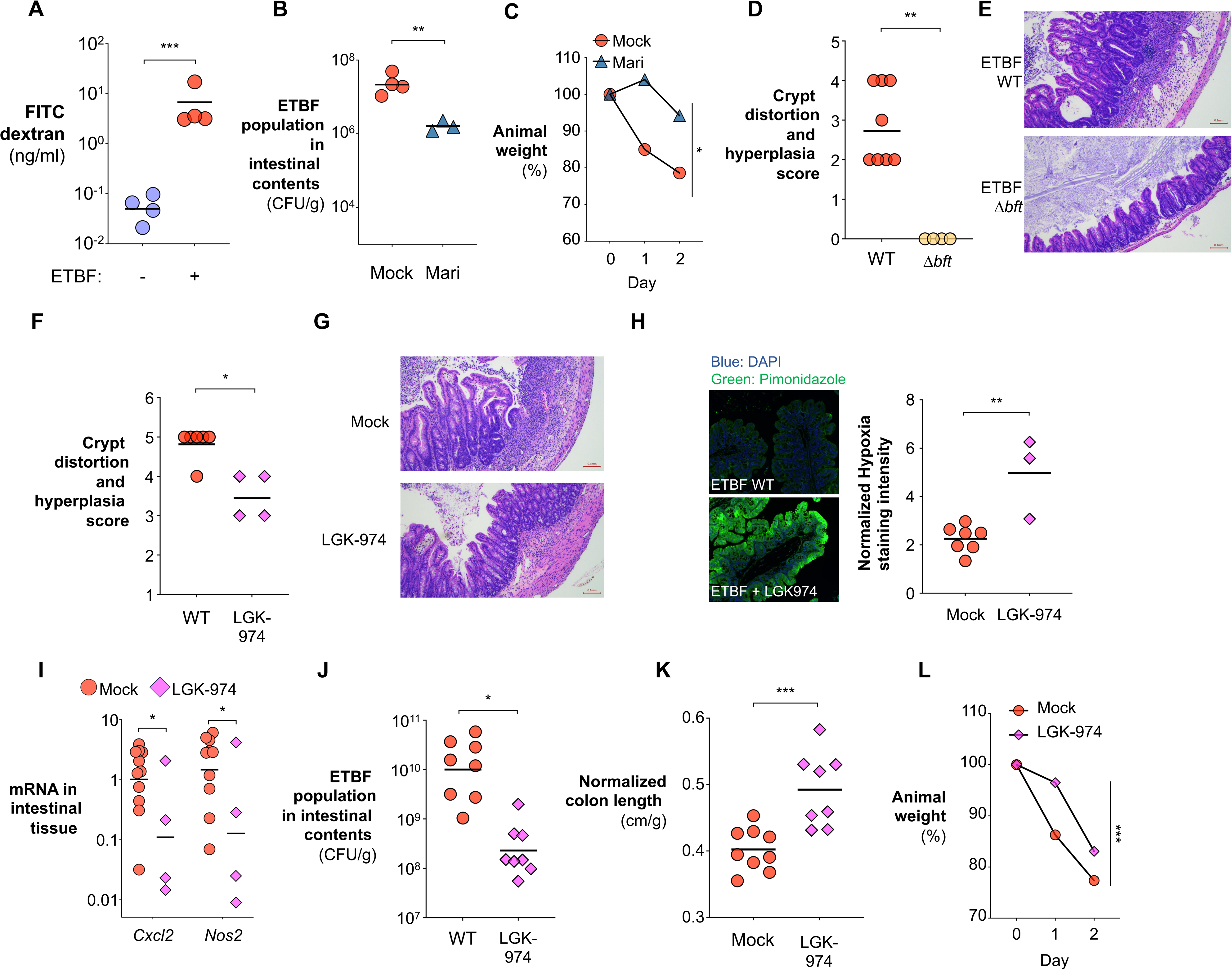
Disruption of epithelial processes promotes ETBF colonization and inflammation. (**A**) C57BL/6 mice were inoculated with either the ETBF wild-type strain or a sham control, followed by intragastric gavage with 4-kDa FITC-dextran. Plasma FITC-dextran levels were measured 4 hours post-gavage to assess intestinal permeability. (**B-C**) ETBF-infected C57BL/6 mice were treated with the MMP inhibitor Marimastat or vehicle control for 3 days. (**B**) Animal weights and (**C**) ETBF burdens in intestinal contents were quantified by plating on selective agar. (**D-E**) C57BL/6 mice were inoculated with either the ETBF wild-type strain or an isogenic Δ*bft* mutant for 7 days. (**D**) Crypt distortion and hyperplasia scores of the cecum tissue. (**E**) Representative H&E-stained images of cecal tissue. (**F-L**) C57BL/6 mice colonized with the ETBF wild-type strain were treated with the Wnt/β-catenin inhibitor LGK-974 or vehicle control for 7 days. (**F**) Crypt distortion and hyperplasia scores in the cecum. (**G**) Representative H&E-stained images of cecal tissue. (**H**) Epithelial oxygenation measured by normalized hypoxia staining. (**I**) ETBF burdens in large intestinal contents quantified by plating. (**J**) Transcript levels of inflammatory cytokines measured by RT-qPCR. (**K**) Normalized colon length. (**L**) Animal weight. Bars represent geometric means. *, *p*<0.05; **, *p*<0.01; ***, *p*<0.001.

E-cadherin proteolysis also leads to the release and nuclear translocation of β-catenin, which activates T-cell factor (TCF)-dependent proliferation, driving epithelial crypt hyperplasia^34, 35, 108^. Indeed, ETBF infection promoted epithelial hyperplasia in a BFT-dependent manner (**Fig. 4D-E**). This excessive division of epithelial cells causes an expansion of undifferentiated, transit-amplifying (TA) cells from their typical location within the crypts to the surface of the epithelium^44^, leading to crypt elongation (**Fig. 4D-E**). Unlike mature colonocytes, TA cells primarily undergo anaerobic glycolysis, which does not contribute to hypoxia^85, 109^. Consequently, colonocyte hyperplasia can promote luminal oxygenation and lactate accumulation.

To assess whether crypt hyperplasia modulates the intestinal nutrient landscape, we inhibited Wnt/β-catenin signaling with LGK-974^110, 111^ during ETBF infection. LGK-974 attenuated ETBF-induced crypt hyperplasia and associated histological distortion (**Fig. 4F-G**). Consistent with a reversal from glycolysis to oxidative metabolism, LGK-974 partially reverted BFT-driven shift towards glycolysis in colonocytes and decreased epithelial oxygenation (**Fig. S4A** and **Fig. 4H**). Importantly, partial normalization of colonocyte metabolism via LGK-974 significantly reduced ETBF loads and alleviated inflammation (**Fig. 4J**), as evidenced by decreased inflammatory cytokine expression, colon shortening, and weight loss (**Fig. 4K-M**). Similar reduction in ETBF colonization was also observed at the peak of infection (Day3, **Fig. S4B**). LGK-974 did not introduce major alterations in overall microbiota composition (**Fig. S4C-G**), indicating that its protective effects are primarily host-mediated. Together, these results indicate that toxin-driven epithelial changes establish a pathological oxidative environment that ETBF exploits for robust colonization.

### BFT-dependent bile acid depletion promotes ETBF colonization

ETBF infection elicits a range of extra-epithelial responses, including a pro-inflammatory cascade that contributes to disease pathogenesis^31^. One hallmark of this cascade is the induction of a pathological T cell program marked by the production of interleukin-17 (IL-17), a cytokine critical for ETBF pathogenesis^27^. IL-17 not only amplifies local inflammation but may also exacerbate metabolic reprogramming in epithelial cells, promoting a shift toward glycolysis^87, 112, 113^ (**Fig. 5A**). However, the mechanisms by which ETBF triggers IL-17 production and how the response feeds back to support bacterial persistence remain incompletely defined.

**Fig. 5.**
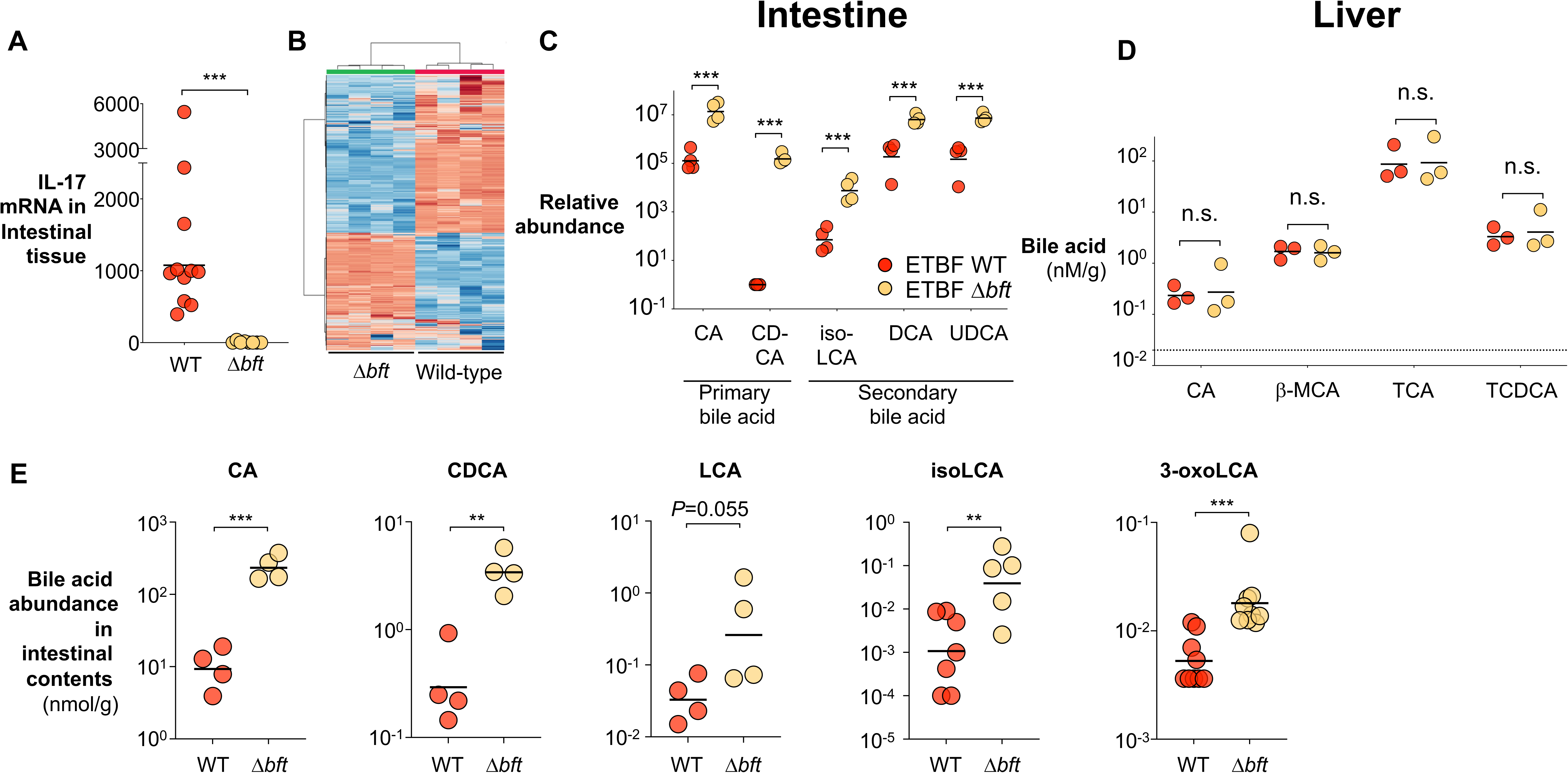
BFT depletes bile acids in the gut. (**A-J**) C57BL/6 mice were colonized with either the ETBF wild-type strain or the isogenic Δ*bft* mutant for 7 days. (**A**) *Il17* mRNA levels in intestinal tissue quantified by RT-qPCR. (**B-C**) Intestinal metabolites were analyzed by UHPLC/MS/MS. (**B**) Heatmap showing differentially abundant metabolites between groups. (**C**) Relative abundance of possible bile acid species in intestinal contents. (**D-E**) Targeted LC-MS/MS quantification of primary bile acids in the (**D**) liver and (**E**) intestinal contents. Bars represent geometric means. n.s., not significant; *, *p*<0.05; **, *p*<0.01; ***, *p*<0.001.

To address this gap, we performed untargeted metabolomic profiling on intestinal contents from ETBF-infected mice (**Fig. 5B**). Remarkably, several primary bile acids, cholic acid (CA) and chenodeoxycholic acid (CDCA), steroidal compounds synthesized in the liver and secreted into the small intestine to aid digestion^114^, were significantly depleted in a BFT-dependent manner (**Fig. 5C**). This loss is unlikely due to reduced biosynthesis, as both hepatic bile acid biosynthesis gene expression and bile acid levels remained unchanged (**Fig. 5D, Fig. S5A-D**).

Similarly, secondary bile acids derived from microbial transformation of these primary bile acids^115^, including deoxycholic acid (DCA), isolithocholic acid (isoLCA), and tauroursodeoxycholic acid (UDCA), were also markedly diminished (**Fig. 5C**). Targeted quantification using LC-MS/MS confirmed BFT-dependent, extensive depletion of both primary and secondary bile acids (**Fig. 5E**). Notably, a subset of secondary bile acids, such as 3-oxolithocholic acid (3-oxoLCA) and isoLCA, potently suppress intestinal T_H_17 differentiation and the resulting IL-17 production^88^ through inhibiting the transcription factor retinoic acid receptor-related orphan receptor γt (RORγt)^88, 116, 117^. This effect likely extends beyond T_H_17 cells, as RORγt also regulates the functions of other IL-17-expressing immune cells, including intraepithelial lymphocytes (IEL), innate lymphocytes, and γδ T cells^118, 119^. As such, the depletion of these secondary bile acids could broadly promote IL-17 signaling through the activation of RORγt-expressing immune cells.

Given the BFT-dependent depletion of 3-oxoLCA and isoLCA observed during ETBF infection (**Fig. 5E**) and the key role of IL-17 in ETBF pathogenesis^120^, we probed whether bile acid depletion promotes ETBF infection by enhancing IL-17 induction. Profiling IL-17 expression in ETBF-infected intestine revealed robust BFT-dependent activation of IL-17-producing, RORγt^+^ immune cells (**Fig. 6A-B**), which include non-T cell IELs (TCR^neg^, RORγt^+^, IL-17^+^), T_H_17 cells (TCRβ^+^, RORγt^+^, IL-17^+^), and γδ T cells (TCRγδ^+^, RORγt^+^, IL-17^+^) (**Fig. 6C-D, Fig. S6A-C**), consistent with previous reports^27, 121, 122^. Importantly, supplementing primary bile acids in drinking water significantly reduced IL-17 production, particularly in non-T cell IELs and γδ T cells, with the latter being critical for ETBF pathogenesis^122^ (**Fig. 6A, C, D, and Fig. S6A-C**).

**Fig. 6.**
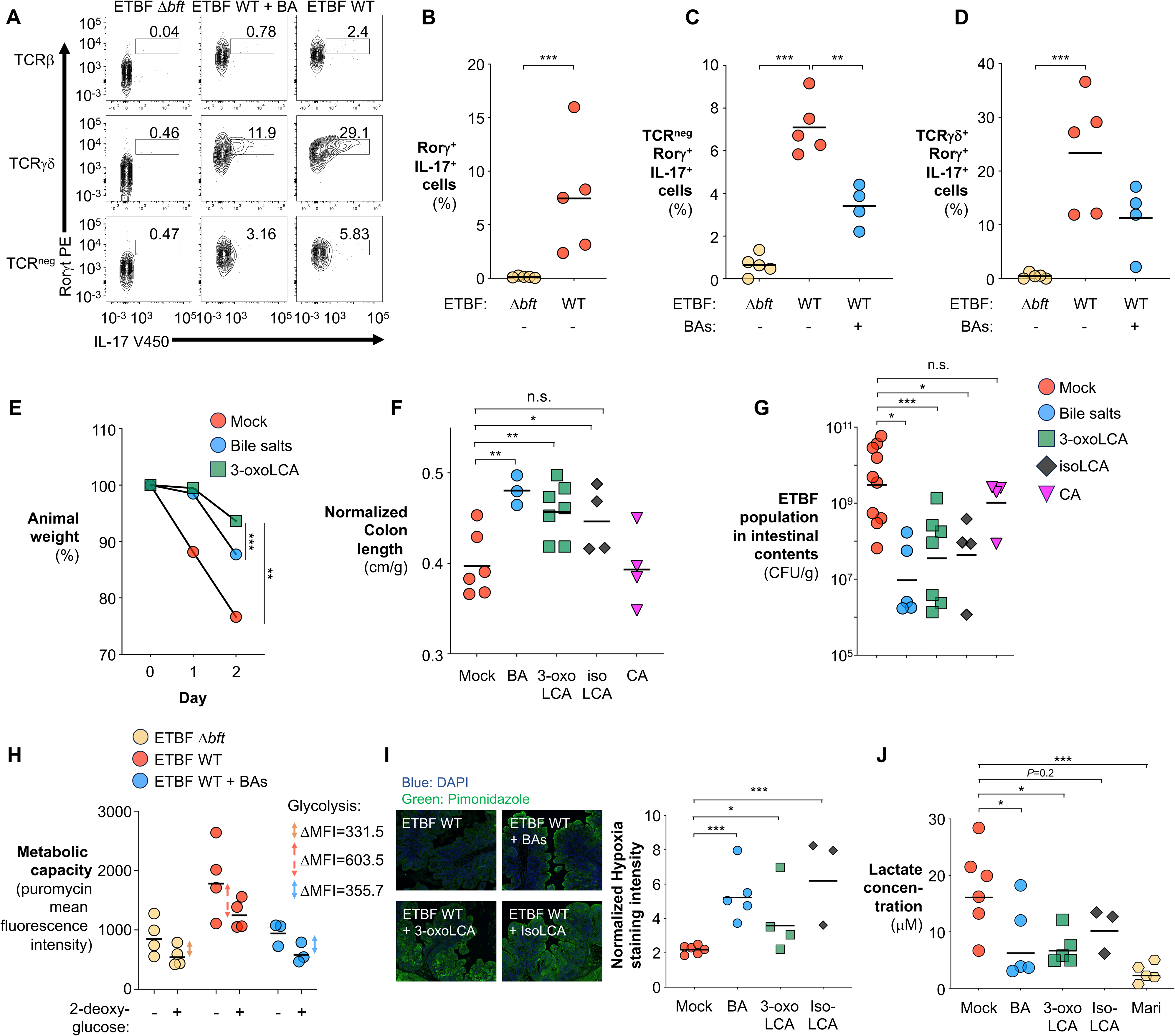
ETBF rewires colonocyte metabolism by impacting bile acid abundance. (**A-D**) Groups of C57BL/6 mice were colonized with wild-type ETBF or the isogenic Δ*bft* mutant and given either vehicle control or primary bile salts in drinking water for 7 days. (**A**) Flow cytometry analysis of IL-17-expressing intestinal lymphocytes. (**B-D**) Quantification of IL-17-producing (**B**) RORgt^+^ immune cells, (**C**) non-T cell IELs, and (**D**) γδ T cells. (**E-J**) Groups of ETBF-colonized C57BL/6 mice were intragastrically administered with the indicated bile acids for 7 days. (**E**) Animal weight. (**F**) Colon length normalized to body weight. (**G**) ETBF abundance in intestinal contents quantified by plating on selective agar. (**H**) Quantification of metabolic activity and glycolytic dependency in gut epithelial cells using SCENITH assay. (**I**) Intestinal oxygenation assessed by hypoxia staining. (**J**) Targeted LC-MS/MS quantification of intestinal lactate levels. Bars represent geometric means. n.s., not significant; *, *p*<0.05; **, *p*<0.01; ***, *p*<0.001.

We next investigated whether bile acids suppress ETBF infection. Under physiological conditions, approximately 95% of the bile acids secreted are recycled back to the liver, while ∼ 5% reaching the colon and undergo microbial transformation into secondary bile acids^123^. To mimic this physiological context, ETBF-infected mice were given a mixture of primary bile acids at a physiological dose (5 μmol/day/kg), matching colonic exposure under homeostasis^123^. Consistent with our hypothesis, supplementation of primary bile acids markedly reduced ETBF-associated animal weight loss, intestinal inflammation, and pathogen colonization (**Fig. 6E-G**). Furthermore, direct supplementation of secondary bile acids, specifically 3-oxoLCA and isoLCA, robustly attenuated ETBF pathogenesis, underscoring secondary bile acid depletion as a critical virulence strategy of ETBF (**Fig. 6E-G, Fig. S6D**). Feature-level clustering analysis of the cecal transcriptome demonstrated distinct molecular signatures associated with supplementation of primary bile acids, 3-oxoLCA, and isoLCA (**Fig. S6E**). Transcriptome analysis further revealed that primary bile acids promoted enrichment of pathways linked to mitochondrial respiration and oxidative phosphorylation, aligning with our observation of restored epithelial metabolic homeostasis (**Fig. S6F-G**). Importantly, supplementation with cholic acid (CA), a primary bile acid that does not yield T_H_17-inhibiting secondary bile acids such as isoLCA and 3-oxoLCA^117^, did not impact ETBF colonization or disease severity (**Fig. 6F-G**). This observation supports the hypothesis that the observed protective effects are specifically mediated by secondary bile acid-dependent immune modulation rather than the general antimicrobial properties of bile acids^124^.

Given that IL-17, the key effector cytokine of the T_H_17 response, has been linked to epithelial adaptation to anaerobic glycolysis^87, 112, 113^, we next examined whether ETBF manipulates epithelial metabolism through a bile acid-IL-17 signaling axis by profiling colonocyte metabolic state using SCENITH assay. Notably, supplementation with primary bile acids suppressed glycolysis in colonocytes (**Fig. 6H**). Furthermore, restoring secondary bile acid levels, either through primary or secondary bile acid supplementation, reduced epithelial oxygenation and luminal lactate accumulation (**Fig. 6I-J**). These treatments also attenuated ETBF-induced T cell differentiation, supporting the idea that pathological T cell response contributes to epithelial metabolic reprogramming (**Fig. S6H**). Together, these findings indicate that ETBF exploits bile acid signaling to reprogram epithelial metabolism, thereby facilitating its colonization of the gut.

### BFT exploits the bile acid recycling pathway to deplete bile acids in the gut

We next investigated the mechanism by which BFT promotes bile acid depletion in the intestinal lumen. As BFT did not alter hepatic bile acid levels or biosynthesis pathway (**Fig. 5D, Fig. S5**), this suggests that processes other than bile acid biosynthesis may be implicated in the observed depletion. Given the majority of the secreted bile acids are returned to the liver via the bile acid recycling pathway, we hypothesized that ETBF may exploit this pathway to reshape the intestinal bile acid landscape^125^. In this recycling pathway, bile acid transporter SLC10A2 uptakes the bile acids into enterocytes, after which fatty acid binding protein 6 (FABP6) shuttles them from the luminal side to the basolateral side of the enterocyte for efflux back to the liver via transporters such as OST-α/β^126^. Transcriptomic profiling of colonic tissues of ETBF-infected mice revealed that BFT upregulated several key components of the bile acid recycling machinery, including Slc10a2, Fabp6, and Ost-α/β (**Fig. 7A**). With bile acid biosynthesis unchanged, enhanced recycling likely reduces the fraction of bile acids that reaches the colon for microbial conversion into immunomodulatory secondary bile acids. Both SLC10A2 and FABP6 proteins are expressed in the large intestine (**Fig. S7A**), supporting the hypothesis that BFT promotes host-mediated bile acid reuptake to deplete secondary bile acids and amplify IL-17-driven inflammation.

**Fig. 7.**
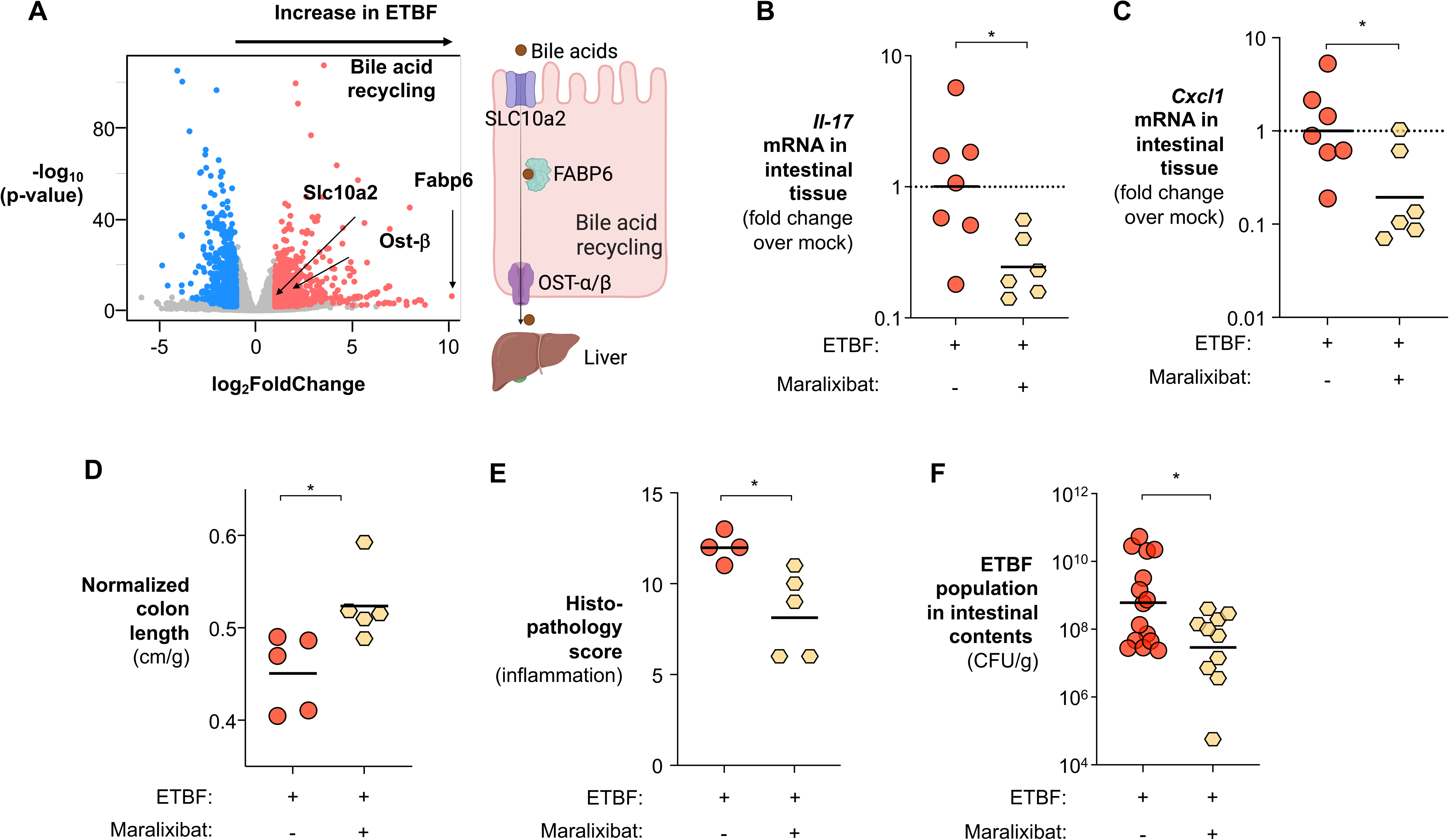
ETBF exploits the bile acid recycling pathway to promote bile acid depletion and gut inflammation. (**A**) C57BL/6 mice were colonized with either wild-type ETBF or the isogenic Δ*bft* mutant for 7 days. The transcriptome of the cecal tissue was profiled by RNA-seq. (**B-F**) ETBF-colonized C57BL/6 mice received a Maralixibat-fortified diet or a control diet for 7 days. (**B**) *Il17* transcript levels in intestinal tissue measured by RT-qPCR. (**C**) *Cxcl1* transcript levels measured by RT-qPCR. (**D**) Colon length normalized to body weight. (**E**) Cumulative histopathology scores assessing cecal inflammation. (**F**) ETBF abundance in intestinal contents quantified by plating on selective media. Bars represent geometric means. *, *p*<0.05.

To further test this, we impeded bile acid recycling using Maralixibat, a selective inhibitor of the primary bile acid transporter SLC10A2^127^. Compared to the vehicle-treated control, Maralixibat treatment restored primary bile acid levels in the large intestine of ETBF-infected mice (**Fig. S7B-C**). Consistent with our hypothesis, this intervention reduced *Il17* and *Cxcl1* transcript levels, markers of IL-17 signaling and inflammation (**Fig. 7B-C**), alleviated colon shortening, and reduced pathogen burden in ETBF-infected animals (**Fig. 7D-F**). Additionally, cecal transcriptome analysis revealed that Maralixibat downregulated genes involved in interferon signaling and γδ T cell activation (**Fig. S7D**), and a broader downregulation of T cell activation pathways was identified by WGCNA (**Fig. S7E**). Importantly, 16S rRNA sequencing of the cecal contents showed augmentation of *Firmicutes*, known critical members of a homeostatic microbiota with no significant changes in alpha diversity, or beta diversity between treatment groups (**Fig. S7F-J**), indicating that the observed effects were unlikely due to shifts in the microbiota. Together, these findings support a model in which BFT manipulates host bile acid recycling to suppress the accumulation of immunomodulatory secondary bile acids in the colon, thereby amplifying IL-17-driven inflammation and promoting ETBF colonization (**Fig. S8**).

## Discussion

To thrive in the competitive environment of the gut, many enteric pathogens deploy virulence factors to reshape the intestinal niche, gaining a competitive advantage over commensal microbes. Pioneering research from the Sears laboratory and others identified *Bacteroides fragilis* toxin (BFT) as the primary virulence factor of ETBF, capable of eliciting profound changes in host physiology. These changes include the cleavage of E-cadherin, induction of the Wnt/β-catenin signaling, and the activation of transcriptional factor STAT3 in gut epithelial cells^82, 83, 128^. In immune cells, BFT also induces robust STAT3 activation, promoting IL-17 induction in subsets such as T_H_17 and δγ-T cells^122^. However, the precise mechanisms by which ETBF benefits from orchestrating these events remain unclear. Here, we show that ETBF reshapes the colonic nutritional landscape by engaging both epithelial and immune cell compartments. BFT drives epithelial hyperplasia and the accumulation of undifferentiated, glycolytic colonocytes. Simultaneously, it activates the bile acid recycling pathway, reducing colonic bile acid levels and derepressing a pathological T cell response. The resulting IL-17 production, in turn, reinforces metabolic reprogramming in colonocytes, creating a feedforward loop that increases oxygen and lactate availability, two metabolites that enhance ETBF colonization (**Fig. S8**).

Our observation that BFT promotes oxygen diffusion into the gut lumen to fuel ETBF colonization has several implications for the pathogenesis of ETBF. First, these observations indicate that ETBF, canonically considered a strict anaerobe, operates an oxidative respiratory metabolism to exploit the altered nutritional landscape for gut colonization. This capability is partly facilitated by enzymes that bridge the two branches of the TCA cycle (e.g., succinyl-CoA synthase SucCD, acetate:succinate CoA transferase ASCT, succinate dehydrogenase Sdh). Together with respiratory lactate dehydrogenases and terminal cytochrome bd oxidase CydAB, ETBF likely runs its electron transport chain more efficiently, generating more ATP and biosynthetic precursors compared to competing gut microbes that rely solely on anaerobic fermentation (**Fig. 1**).

This respiratory capability may explain previous findings that *B. fragilis* can utilize nanomolar oxygen (nanaerobe^66, 73^) and appears to be better adapted than other *Bacteroidetes* to thrive within the mucosal environment. Of note, our results suggest that ETBF can grow robustly in the presence of micromolar oxygen, suggesting that this pathogenic strain may have metabolically adapted to the oxidative, inflamed gut. We do not know why ETBF exhibits higher oxygen tolerance than the non-enterotoxigenic *B. fragilis*. It’s plausible that genes on the *B. fragilis* pathogenicity island (BFPAI), acquired through horizontal gene transfer, may confer enhanced resistance to oxygen and reactive oxygen species (ROS) via an uncharacterized mechanism ^129, 130^. Further research is needed to uncover the specific mechanisms involved.

While ETBF’s capacity for oxidative metabolism improves its fitness during inflammation, it retains the ability to ferment complex glycans^131^. This metabolic flexibility is consistent with the significant, but relatively modest *in vivo* phenotype observed when key components of ETBF’s oxidative metabolism were disrupted (**Fig. 2**). These effects may have been further obscured by a key limitation of our study: the requirement to use antibiotics to permit consistent, robust ETBF colonization^27, 35^. Antibiotic treatment depletes resident microbiota, potentially opening nutrient niches that are normally occupied. This ecological disruption could also liberate alternative nutrient sources, thereby masking the specific contributions of oxygen and lactate, metabolites elevated through BFT-mediated host metabolic reprogramming, to ETBF fitness. Although the current murine model enabled us to dissect the role of oxidative metabolism in ETBF pathogenesis, future development of antibiotic-free colonization models may offer a more physiologically relevant framework to uncover the full spectrum of metabolic strategies that ETBF employs to colonize the inflamed gut.

Beyond supporting the oxidative metabolism in ETBF, increased oxygen and ROS could damage oxygen-sensitive enzymes in anaerobic commensals^99^. This metabolic disruption may force these microbes to divert pyruvate toward lactate production^46, 100^. The microbially-derived lactate, often in the D-enantiomer form, could contribute to the increase in the total lactate pool. Consistently, ETBF codes for dehydrogenases capable of utilizing both L- and D-lactate (LutABC & Ldh2 and Ldh1, respectively), positioning it to exploit the expanded lactate pool. Moreover, increased oxygen and ROS levels are associated with the depletion of less aerotolerant species that may otherwise compete with ETBF for similar nutrients or by deploying a bactericidal type VI secretion system^52, 97, 129, 132, 133^. Thus, by sculpting an oxidative niche, ETBF both fuels its own growth and suppresses its microbial competitors. Importantly, this distinct metabolic program could be potentially leveraged to selectively target and remove ETBF. For example, we previously exploited the molybdenum cofactor dependency of *Enterobacteriaceae* and used tungstate to selectively inhibit their inflammatory expansion^134^. Similarly, targeting ETBF’s dependence on metabolically reprogrammed colonocytes could limit its persistence in the gut (**Fig. 3**).

The bile acid–IL-17 signaling axis also offers a potential therapeutic entry point. However, given the broad physiological roles of both primary and secondary bile acids, including regulation of intestinal barrier integrity^135^, wound healing response^136^, immune system functions^88^, and gut microbiota composition^115^, strategies to restore bile acid homeostasis need to be approached with caution. These broad impacts of bile acids on intestinal physiology likely contribute to the modest phenotypic impact observed with Maralixibat, a selective inhibitor of the bile acid recycling pathway (**Fig. 7**). For example, excessive inhibition of bile acid reabsorption could lead to accumulation of deoxycholic acid, a secondary bile acid known to impair colonic epithelial wound healing^136, 137^. This could paradoxically exacerbate ETBF infection by further compromising epithelial integrity and enhancing bacterial access to the lamina propria. These findings underscore the need to explore combination strategies, such as pairing Maralixibat with agents that promote epithelial repair, to optimize therapeutic efficacy against ETBF infection while minimizing unintended physiological disruptions.

In summary, this work uncovers a previously underappreciated link between bile acid metabolism, IL-17 response, and colonocyte metabolism. We establish a mechanistic crosstalk between the enteric pathogen ETBF and colonocyte metabolism in the context of colonic inflammation, revealing how toxin-driven metabolic rewiring of colonocytes creates an oxidative niche that supports selective anaerobic pathogen colonization (**Fig. S8**).

## STAR METHODS

### Bacterial strains, growth conditions and ATP assay

Enterotoxigenic *Bacteroides fragilis* (ETBF) strain 86-5443-2-2 was grown anaerobically (90% N_2_, 5% CO_2_, 5% H_2_; vinyl anaerobic chamber, Coy Lab) in brain-heart-infusion supplemented (BHIS) media (0.8% brain heart infusion from solid, 0.5% peptic digest of animal tissue, 1.6% pancreatic digest of casein, 0.5% sodium chloride, 0.2% glucose, 0.25% disodium hydrogen phosphate, 0.005% haemin, 0.0001% /vitamin K, pH 7.4) or BHIS plates (BHIS broth, 15 g/L agar) containing 50 μg/mL gentamycin (Gen) for 2 days at 37 °C. When appropriate, agar plates and media were supplemented with 50 μg/ml kanamycin (Kan), 25 μg/ml erythromycin (Erm), 25 μg/ml clindamycin (Clin) or 200 μg/ml 5-fluoro-2′-deoxyuridine (FUdR).

For microaerobic growth, single colonies of indicated strains were cultured anaerobically for 24 hours in BHIS before subculturing into modified semi-defined media (SDM)^138^ (1.5 g/L KH_2_PO_4_, 0.5 g/L NH_4_SO_4_, 0.9 g/L NaCl, 150 mg/L L-methionine, 5 μg/L vitamin B_12_, 1 mg/L resazurin, 1 g/L tryptone, 0.2% NaHCO_3_, 0.005% protoporphyrin IX) supplemented with 1 mM lactate. Culture were inoculated at the density of (2.5x10^4^ CFU) into an open, micro-anaerobic atmosphere 1-10 μM O_2_ (0.1-1% O_2_, 5% CO_2_, 2.5% H_2_; In vitro Hypoxia system, Coy Lab) and incubated at 37 °C for up to 72 hours.

For ATP assay, single colonies were cultured in SDM anaerobically before subculturing into lactate supplemented SDM in microaerobic atmosphere (0.1 % O_2_) for 16 hours. Cells were washed and resuspended SDM. 2.5x10^6^-2.5x10^7^ cells were used for assaying ATP concentration using an ATP determination kit (BacTiter-Glo Microbial Cell Viability Assay kit, Promega). Sample luminescence was read using white-well plates in a BioTek H1 plate reader and compared to a standard curve of known ATP concentrations.

*E. coli* strains were routinely cultured in LB broth (10 g/L tryptone, 5 g/L yeast extract, 10 g/L sodium chloride) or on LB plates (LB broth, 15 g/L agar) at 37 °C supplemented with 100 µg/mL carbenicillin.

### Plasmids

All the primers and plasmids used in this study are listed in the Key Resource Table. Suicide plasmids were routinely propagated in *E. coli* S17 λ*pir*. The flanking regions of ETBF strain 086-54443-2-2 *cydAB, lutABC, ldh1, ldh2, asct, su<u>c</u>AB, su<u>c</u>CD* were amplified and assembled into pETBF-Exchange-*tdk* (pEET) using the Gibson Assembly Cloning Kit (New England Biolab, Boston) to give rise to pRF237, pRF231, pML102, pML108, pML191, pRF317, and pRF247, respectively. The ETBF 11*bft* strain is a generous gift from Dr. Cynthia Sears^31^.

### Construction of mutants by allelic exchange

All bacterial mutant strains constructed using the method below are listed in the Key Resource Table. For ETBF mutants, suicide plasmid pEET containing the flanking regions of genes of interest was conjugated using *E. coli* S17-1 λ*pir* as the conjugative donor strain into the ETBF. Exconjugants with suicide plasmid integrated into the recipient chromosome were selected on BHIS plates supplemented with appropriate antibiotics. 5-fluoro-2-deoxy-uridine (FudR, 200 µg/mL in BHIS) plates were used to select for the second crossover event.

### Phylogenetic analysis of genes involved in oxidative central metabolism

RefSeq genome annotations were downloaded using the NCBI Datasets command line tool^139^ (v16.40.1) on April 8^th^, 2025. Complete assemblies for *Bacteroides fragilis, Phocaeicola vulgatus, Bacteroides uniformis, Bacteroides ovatus, Bacteroides thetaiotaomicron, Bacteroides caccae, Bacteroides salyersiae, Bacteroides xylanisolvens, Bacteroides cellulosilyticus,* and *Phocaeicola dorei* were downloaded, as well as scaffold assemblies for *Hoylesella oralis, Bacteroides acidifaciens, and Bacteroides nordii*, which were used due to a lack of complete assemblies. Only the latest assembly version was used, and metagenome assembled genomes, assemblies from large, multi-isolate projects, and atypical assemblies were excluded. Annotated protein sequences were used as input for Orthofinder^140^ (v3.0.1b1), which produced the phylogeny and the presence/absence calls for each orthogroup. A phylogeny for the species was using ggtree^141^ (v3.16.0).

### Animal Experiments

All experiments were conducted in accordance with the policies of the Institutional Animal Care and Use Committee at Vanderbilt University Medical Center. C57BL/6J wild-type (cat# 000664), were obtained from Jackson Laboratory (Bar Harbor) and housed in sterile cages under specific pathogen-free conditions on a 12-hour light cycle, with *ad libitum* access to food and sterile water at Vanderbilt University Medical Center.

Three-week-old male and female mice were randomly assigned into treatment groups before the experiment. Streptomycin (5 mg/ml) and clindamycin (0.1 mg/ml) were administered in drinking water for 2 days, followed by 3 days of clindamycin only. On day 6 mice were then inoculated with 1 x 10^9^ CFU of the indicated enterotoxigenic *Bacteroides fragilis* strains or remained uninfected.

For the tumorigenesis model, three-week-old *Apc*^Min^ male and female mice, together with their littermate controls were randomly assigned to treatment groups before the experiment. Streptomycin (5 mg/ml) and clindamycin (0.1 mg/ml) were administered in drinking water for 2 days, followed by 3 days of clindamycin only. On day 6 mice were then inoculated with 1 x 10^9^ CFU of the indicated enterotoxigenic *Bacteroides fragilis* strains or remained uninfected, followed by a single dose of tributyrin treatment. 12 weeks post infection, animals were euthanized, colonic tissue fixed with 4% paraformaldehyde, and tumor incidence was quantified.

For all experiments, mice were humanely euthanized at the indicated time points. After euthanasia, liver, cecal, and colonic tissue were collected, flash-frozen, and stored at -80 °C for subsequent mRNA analysis or fixed in 10% formalin for histopathological analysis. For untargeted metabolomics, cecal contents were collected and flash frozen in liquid nitrogen, stored in -80 °C until metabolites were extracted. For culture-dependent quantification of bacterial load, colonic and cecal contents were harvested in sterile PBS, and the load of enterotoxigenic *Bacteroides fragilis* (ETBF) was quantified by plating serial-diluted intestinal contents on selective agar.

### Hybrid Selection-RNAseq

Groups of C57BL/6 were challenged with either ETBF wild-type strain or the isogenic Δ*bft* as described above. After euthanasia, cecal samples was lysed immediately after collection in 500µl buffer (0.2M of NaCl and 20mM of EDTA), 210µl of 20% SDS, and 500µl of phenol, chloroform and isoamyl alcohol mixture using lysing Matrix B. Total RNA was extracted and purified from three sites per mouse (intestinal tissue, mucus layer, and luminal contents) across eight mice (24 biological samples in total) using the RNeasy Mini Kit.

RNA was used to generate Illumina cDNA libraries using a modified version of the RNAtag-seq protocol^142, 143^. Briefly, RNA was fragmented, depleted of genomic DNA, dephosphorylated, and ligated to DNA adapters carrying 5′-AN₈-3′ barcodes of known sequence with a 5′ phosphate and a 3′ blocking group. To ensure sufficient input materials for hybrid capture, libraries were generated using 2.25 ug of total RNA per sample split across 9 technical replicates processed in parallel. Barcoded RNAs from four biological samples were combined in each pool. Pooled RNA underwent rRNA depletion using the Human RiboCop rRNA Depletion Kit (Lexogen, Cat. No. 144), followed by reverse transcription using a primer targeting the constant region of the barcoded adapter, with 3′ adapter addition via template switching (SMARTScribe, Clontech)^144^. cDNA was enriched by PCR for 14 cycles using primers targeting the constant regions of the adaptors with attached Illumina P5/P7 sequences carrying unique indexes per pool. cDNAs from replicate pools generated in parallel were combined and concentrated and used as input for enrichment. The yield of library pool after 14 PCR cycles ranged from 350 (mucus, ETBF Δ*bft*) to 1,600 ng (lumen, ETBF WT). These amplified indexed libraries containing barcoded samples were subjected to hybrid capture enrichment as described below.

Enrichment of ETBF sequences using a custom capture panel and v1 hybridization and wash reagents (Twist Biosciences) was performed in 6 separate capture reactions, each containing approximately 250 ng pre-capture RNAtag library pool. We followed the manufacturer’s instructions except that we added 5 ug mouse Cot-1 DNA (Thermo Fisher) instead of the human blocker solution to the hybridization and used HiFi HotStart ReadyMix (Roche) for PCR amplification of the catch. To empirically determine the minimum cycle number to generate approximately equal quantities of PCR product despite different amounts of ETBF in the three intestinal sites (highest in lumen, lowest in tissue) and ETBF WT (higher) vs. ETBF Δ*bft* (lower), we performed scaled-down 10 ul PCR reactions from 2 ul capture-bead slurry with 8, 12 or 16 cycles. The number of PCR cycles necessary to generate ∼100 fmol of each capture library from 25 ul bead slurry ranged from 5 (ETBF WT in lumen) to 11 (tissue).

Pre-capture RNAtag pools were sequenced on two Illumina NovaSeq S4 lanes to an average depth of 9.4x10^8^ read pairs per sample. Capture libraries were sequenced on one lane to an average depth of 4.8x10^8^ read pairs per sample. The sequencing reads generated are available at the NCBI BioProject Database under accession number PRJNA1269620.

### Analysis of Hybrid Selection-RNAseq data

Sequencing reads from each sample in a pool were demultiplexed based on their associated barcode sequence using custom scripts (https://github.com/broadinstitute/split_merge_pl). Up to 1 mismatch in the barcode was allowed provided it did not make assignment of the read to a different barcode possible. Barcode sequences were removed from the first read as were terminal G’s from the second read that may have been added by SMARTScribe during template switching.

*Bacteroides fragilis*-derived reads were aligned to a the GCF 023702735.1 ASM2370273v1 reference sequence using BWA^145^ and read counts were assigned to RefSeq annotated genes and other genomic features using custom scripts (https://github.com/broadinstitute/BactRNASeqCount). Mouse-derived reads were aligned to *Mus musculus* Ensembl sequence GRCm38r94p6 mm10 using bbmap_37.10 (https://jgi.doe.gov/data-and-tools/bbtools/) and read counts were assigned to annotated transcripts using Salmon_0.8.2^146^.

Differential expression analysis was conducted with DESeq2^147^ and/or edgeR^148^. Visualization of raw sequencing data and coverage plots in the context of genome sequences and gene annotations was conducted using GenomeView^149^.

### Targeted quantification of mRNA levels in intestinal tissue and contents

Colonic or cecal tissue was homogenized in a bead beater (Precellys 24 Touch, Bertin Technologies) and RNA was extracted using TRI reagent (Molecular Research Center, Cincinnati). DNA contamination was removed using the Turbo DNA-free Kit (Ambion, USA) per the manufacturer’s recommendations. cDNA was generated by SuperScript VILO cDNA Synthesis Kit (Thermo Fisher, USA). Real-time PCR was performed using PowerUp SYBR Green Master Mix (Applied Biosystem, USA), data were acquired in a CFX Maestro 2.3 (Bio-Rad, USA). Target gene transcription of each sample was normalized to *Gapdh* mRNA levels.

### Western blot analysis of FABP6 and SLC10A2 expression in intestinal tissue

Colonic proteins were extracted using RIPA lysis buffer. For each 5 mg of tissue, 600 µL of ice-cold lysis buffer were added, and samples were homogenized using a bead beater (Precellys 24 Touch, Bertin Technologies). Lysates were agitated for 2 hours at 4 °C and then centrifuged at 16,000 g for 20 minutes at 4 °C. Supernatants were collected, and protein concentrations were determined by measuring absorbance at 280 nm using an Epoch Microplate Spectrophotometer (BioTek Instruments).

Samples were boiled at 95 °C for 1 minute, and 50 μg of total protein per sample were resolved on 4–20% TGX precast gels (Bio-Rad). Proteins were transferred onto polyvinylidene fluoride (PVDF) membranes via wet transfer (Bio-Rad). Membranes were blocked in TBS containing 3% non-fat dry milk and 0.1% Tween-20, then incubated overnight at 4 °C with primary antibodies against FABP6 and SLC10A2. After washing, membranes were incubated with appropriate IRDye-conjugated secondary antibodies for 1 hour at room temperature and scanned using an Odyssey Western Blot Imagers (LICORbio). Relative expression levels were quantified using ImageJ, with uniform adjustment of brightness levels across samples.

### Colonocyte Isolation

Colon and cecum of ETBF-infected animals were removed and kept cold in RPMI + 10% FBS. Luminal contents were flushed out with HBSS + 5% FBS + 2mM EDTA. The colon and cecum were then cut longitudinally and into smaller pieces, placed in HBSS medium containing 5% FBS + 2mM EDTA + DTT (1.5 ml/ml) and shake at 37 °C for 50 minutes at 150 rpm. After the incubation, the supernatant and the tissue were separated, with the supernatant filtered and centrifuged for 5 minutes at 1200 rpm at 4 °C. The resulting pellet was resuspended in HBSS + 5% FBS + 2mM EDTA and kept on ice. The tissue was digested in HBSS (contains Ca^2+^ and Mg^2+^) supplemented with collagenase (1.5 mg/ml) and DNase 1 (1000X). After shaking for 25 minutes at 100 rpm, HBSS + 5% FBS + 2mM EDTA was added to stop the digestion, the remaining tissue was filtered out using cell strainer, and the digested cells were centrifuged for 5 minutes at 1200 rpm at 4 °C. After centrifugation, the two populations (from supernatant and tissue) of cells were combined and further purified using Percoll gradient centrifugation. Cells at the interface of the 40% and 70% layer were collected, centrifuged for 5 minutes at 1200 rpm at 4 °C, washed using RPMI + 10% FBS, and finally resuspended in 500 μL of the same media. Cell count and viability were quantified using Countess 3 cell counter (Thermo Fisher Scientific).

### SCENITH assay

Cells were seeded at 1 x 10^6^ cells/ml of RPMI + 10% FBS in 96-well plates and incubated in a 37 °C CO_2_ incubator for 1 hour. Wells were treated with DMSO Control, 2-Deoxy-D-Glucose (DG, final concentration 250 mM) or Oligomycin (Oligo, final concentration 1 μM). Puromycin (final concentration 10 μg/ml) was added, and the cells were returned to the incubator for 20 minutes. After puromycin treatment, cells were washed in cold PBS and stained with a 1:100 cocktail mix of anti-CD326-APC, CD45 RED Fluo 710, TCR gamma/delta Fluo 450 antibodies, for 30 minutes at 4°C in PBS, 5% FCS and 2 mM EDTA (FACS wash buffer). After washing with cold PBS, ghost viability dye (BV510) 1:100 in FACS buffer was added, and cells were incubated at 4 °C for 20 minutes in the dark. After incubation, cells were washed in FACS buffer and fixed and permeabilized using FOXP3 fixation and permeabilization buffer (Thermofisher eBioscience) following manufacturer’s instructions. Intracellular staining of puromycin was performed by incubating cells for 1 hour at RT with anti-Puromycin antibody (PE) 1:100 in permeabilization buffer. After washing, cells were resuspended in 250ml FACS buffer (2% FBS + 2mM EDTA in PBS) and transferred in FACS tubes and data were acquired using Fortessa (BD Biosciences). Flow cytometry data were analyzed using FlowJo (BD Biosciences).

### IL-17-producing immune cell profiling

100,000 to 500,000 cells per well were plated 200 μl/well of stimulation buffer (RPMI + 10 % FBS, 0.67 μl/ml GolgiStop protein transporter inhibitor, 500 μg/ml ionomycin, 25 μg/ml phorbol 12-myristate 13-acetate (PMA)) in 96-well plates, and incubated in CO_2_ incubator at 37 °C for 4 hours. Cells were stained with 1:100 TCR γ/δ biotinylated antibodies in FACS buffer (2% FBS + 2 mM EDTA in PBS), and incubated 25 minutes at 4°C. After incubation, cells were washed and incubated with a 1:100 cocktail mix of Streptavidin PE610, CD45 PECy7, TCRβ APC Cy7 for 25 minutes at 4°C. Cells were washed with cold PBS and ghost viability dye (BV510) 1:100 in FACS buffer was added and cell were incubated at 4 °C for 20 minutes in the dark. After incubation, cells were washed in FACS buffer, fixed and permeabilized using FOXP3 fixation and permeabilization buffer (Thermofisher eBioscience). Cells were washed with 150 μl/well of Perm-Wash buffer (Thermofisher eBioscience), and incubated in the dark at RT for 40 minutes with an intracellular antibody cocktail in Perm-Wash buffer. After incubation, cells were washed with Perm-Wash buffer, followed by a second wash in FACS buffer. Cells were finally resuspended in 200 μl FACS buffer and transferred to FACS tubes to be acquired in a FACS machine. Data were acquired using Fortessa (BD Biosciences), and flow cytometry data were analyzed using FlowJo (BD Biosciences).

### Histopathology

Cecal and colonic tissue were fixed in phosphate-buffered formalin for 48 h and embedded in paraffin. Sections were stained with hematoxylin and eosin. Stained sections were blinded and evaluated according to the criteria listed in the **Supplementary Table S2**.

### Hypoxia staining

ETBF-infected mice with indicated treatment were peritoneally injected with 60 mg/kg pimonidazole (Hypoxyprobe) 30 minutes prior to euthanasia. After euthanasia, samples are collected as described above and fixed in 10% formalin. Tissues were embedded, sectioned and mounted on glass slides. Slides of cecal and colonic tissue were deparaffinized and incubated for 10 minutes at 37 °C with 200 ml of proteinase K (New England Biolabs) solution in TE buffer (10 mM Tris, 1 mM EDTA, pH=8.0). The slides were washed in PBS and blocked for 1 hour at 4 °C with 150 ml of M.O.M mouse IgG Blocking reagent (Vector Laboratories). Samples were incubated with 100 μl of α-pimonidazole rabbit antisera (Hypoxyprobe) at a 1:100 ratio overnight at 4 °C. Samples were washed in PBS and incubated with 100 μl of Alexa-488 conjugated goat anti-rabbit secondary antibody (Cell Signaling, 1:150) for 90 minutes at RT. After washing with PBS, slides were counterstained with Hoechst for 5 minutes, followed by additional PBS washes and mounting. Tiled fluorescence images (5 × 5 fields, 15% overlap) were acquired with Nikon ECLIPSE Ti-2 fluorescence microscope with a 20x objective and the Perfect Focus System enabled. Imaging stitching was performed in real time using NIS-Elements (version 6.10.01). Regions of interest (ROI) were selected manually to exclude well-oxygenated lamina propria. The intensity of pimonidazole staining within each ROI was normalized to the Hochest signal in the same region to yield normalized Hypoxia staining intensity.

### Quantification of lactate

Colon and cecal contents of ETBF-infected mice were collected in sterile PBS. Contents were weighed, diluted with 1x PBS to a final density of 40 mg/mL, and homogenized by gentle vortexing at 5 °C. Insoluble debris was removed by centrifugation at 3,000 xg for 10 minutes at 5 °C. The supernatants were collected and stored at -20 °C until the day of analysis.

Lactate and a stable isotope-labeled internal standard lactate-*^13^C*_3_ (Cambridge Isotopes) were derivatized with the reagent dansyl hydrazine and the carboxyl activating agent 1-Ethyl-3- (3-dimethylaminopropyl)carbodiimide (EDC) to their corresponding dansyl hydrazone derivatives in 250 mM aqueous sodium phosphate pH 4 buffer containing 40% (*v/v*) acetonitrile. Briefly, 20 μL aliquots of thawed extracts were spiked with 5 nmol lactate-*d*_3_, diluted 1:7 with the derivatization buffer described above, and derivatized at room temperature with the addition of dansyl hydrazine (25 μL x 50 mg/mL in acetonitrile) and EDC (25 μL x 150 mg/mL in water). After one hour at room temperature, 25 μL water containing 5% (*v/v*) TFA was added to quench the reactions. Following centrifugation at 10,000 x *g* for 30 min at 5 °C, quenched reaction mixtures were transferred to 2-mL autosampler vials equipped with low-volume polypropylene inserts and Teflon-lined rubber septa. The sample injection volume was 10 μL. Calibration standards were prepared in 1x PBS and derivatized in the same manner. LC-MS/MS analysis was performed using a Thermo TSQ Quantum triple-stage quadrupole mass spectrometer interfaced to a Waters Acquity UPLC system. The mass spectrometer was operated in positive ion mode, with detection by multiple reaction monitoring (MRM) using the following transitions: Lactate, 338 → 170, CE 27; Lactate-^13^C_3_, 341 → 170, CE 27. The following analyzer and ESI source parameters were used for the detection of analyte and internal standard: Q1 peak width 0.7 Da (FWHM), Q3 scan width 2 Da, spray voltage 4 kV, capillary temperature 300 °C, tube lens 120, N_2_ sheath gas 50; N_2_ auxiliary gas 10; in-source CID 12. Data acquisition and quantitative spectral analysis were done using Thermo Xcalibur version 2.0.7 SP1 and Thermo LCQuan version 2.7, respectively. Calibration curves were constructed by plotting peak areas against analyte concentrations for a series of ten calibration standards, ranging from 0.10 to 1000 total nmol lactate. A weighting factor of 1/C^2^ was applied in the linear least-squares regression analysis to maintain homogeneity of variance across the concentration range. A Waters Acquity BEH C18 reverse phase analytical column (2.1 x 100 mm, 1.7 μm) was used for all chromatographic separations. Mobile phases were made up of 0.2 % HCOOH + 10 mM NH_4_^+^ Formate in (A) water and in (B) acetonitrile/methanol (9:1). Gradient conditions were as follows: 0–1.0 min, B = 5 %; 1–8 min, B = 5–100 %; 8–9.5 min, B = 100 %; 9.5–10 min, B = 100–5 %; 10–15 min, B = 5 %. The flow rate was maintained at 300 μL/min, and the total chromatographic run time was 15 min. A software-controlled divert valve was used to transfer the LC eluent from 0 to 3 min and from 5.5 to 15 min of each chromatographic cycle to waste.

### Bile acid quantification

Bile acids (BAs) were measured as their Girard P hydrazide derivatives^150^. Briefly, to 25 μL of homogenate, a mixture of deuterium-labeled bile acid internal standards was added (5 μL x 25 μg/mL of each: cholate-d_4_, deoxycholate-d_4_, glycocholate-d_4_, chenodeoxycholate-d_4_, ursodeoxycholate-d_4_, and lithocholate-d_4_; all from CDN Isotopes). Spiked samples were then extracted with 750 μL of a solution of methanol/methyl tert-butyl ether/chloroform (1.3:1:1 v/v/v). Following centrifugation at 10,000 x g, the supernatants were removed and evaporated under a gentle stream of nitrogen gas. Residues were reconstituted in 100μL pyridine hydrochloride buffer (300 mM, pH 4.5) before adding 25 μL x 50 mg/mL Girard P reagent in acetonitrile/water (1:1) and 25 μL x 150 mg/mL of the carboxyl activating agent 1-Ethyl- 3-(3-dimethylaminopropyl)carbodiimide (EDC) in water to convert BAs to their corresponding Girard P hydrazides. After shaking overnight at 4 °C, reactions were quenched with 25 μL x 5 % (v/v) TFA in water. Quenched reactions were centrifuged at 18,000 x g, then transferred to 2 mL autosampler vials equipped with low-volume polypropylene inserts and Teflon-lined rubber septa. The sample injection volume was 10 μL. BA calibration standards were prepared in water and derivatized in the same manner. Liquid chromatography-high resolution mass spectrometry (LC-HRMS) analysis was performed using a Thermo Q Exactive HF hybrid quadrupole/orbitrap high resolution mass spectrometer interfaced to a Vanquish Horizon HPLC system (Thermo Fisher). High resolution mass spectra were acquired in positive ion mode over a precursor ion scan range of m/z 300 to 800 at a resolving power of 60,000 using the following ESI source parameters: spray voltage 4 kV; capillary temperature 300°C; HESI temperature 100 °C; s-lens 95; N2 sheath gas 40; N2 auxiliary gas 10. Extracted ion chromatograms were constructed for each BA based on the following [M+H]+ exact masses and a mass tolerance of +/- 5 ppm: CA, 542.3588; CA-d4, 546.3840; DCA and CDCA, 526.3639; DCA-d4 and CDCA-d4, 530.3890; LCA and iso-LCA, 510.3690; LCA-d4, 514.3941; 3-oxo-LCA, 508.3534. Data acquisition and quantitative spectral analysis were done using Thermo Xcalibur version 4.1.31.9 and Thermo LCQuan version 2.7, respectively. Calibration curves were constructed by plotting peak areas against analyte concentrations for a series of eleven calibration standards, ranging from 0.15 to 1500 total pmol of each BA. A weighting factor of 1/C2 was applied in the linear least-squares regression analysis to maintain homogeneity of variance across the concentration range. An Acquity HSS C18 reverse phase analytical column (2.1 x 150 mm, 1.7 μm, Waters, Milford, MA) was used for all chromatographic separations. Mobile phases were made up of 0.2 % formic acid + 10 mM ammonium formate in (A) water/methanol/acetonitrile (8:1:1) and in (B) methanol/acetonitrile (4:1). Gradient conditions were as follows: 0–1.0 min, B = 10 %; 1–15 min, B = 10–100 %; 15–17.5 min, B = 100 %; 17.5–18 min, B = 100–10 %; 18–22 min, B = 10 %. The flow rate was maintained at 300 μL/min, and the total chromatographic run time was 22 min. A software-controlled divert valve was used to transfer the LC eluent from 0 to 5 min of each chromatographic cycle to waste.

### Untargeted metabolomics

Samples were stored at -80°C until analyzed via liquid chromatography-high resolution tandem mass spectrometry (LC-HRMS and LC-HRMS/MS)- based metabolomics in the Vanderbilt Center for Innovative Technology (CIT) using previously described methods (^151–153^. Briefly, frozen mouse intestinal content samples (n=8, 4 biological replicates for each sample group) were lysed in 1 ml ice-cold lysis buffer (1:1:2, v:v:v, acetonitrile: methanol: ammonium bicarbonate 0.1M - pH 8.0) and sonicated individually using a probe tip sonicator at 50% power (10 pulses). Homogenized samples were normalized by weight to an equal amount per sample. Proteins were precipitated from individual samples by adding 800 µL of ice-cold methanol, followed by overnight incubation at -80°C. Precipitated proteins were pelleted by centrifugation (15k rpm, 15 min), and metabolite extracts were dried in vacuo.

Individual extracts were reconstituted in 100 µl of acetonitrile/water (3:97, v:v) with 0.1% formic acid containing isotopically-labeled carnitine-D9, tryptophan-D3, valine-D8, and inosine-4N15, and centrifuged for 5 min at 10,000 rpm to remove insoluble material. A pooled quality control (pooled QC) sample was prepared by pooling equal volumes of individual samples. The pooled QC was used for column conditioning (8 injections prior to sample analysis), retention time alignment and to assess mass spectrometry instrument reproducibility throughout the sample set.

Global, untargeted mass spectrometry analyses were performed on a high-resolution Q-Exactive HF hybrid quadrupole-Orbitrap mass spectrometer (Thermo Fisher Scientific, Bremen, Germany) equipped with a Vanquish UHPLC binary system (Thermo Fisher Scientific, Bremen, Germany). Extracts (5 μL injection volume) were separated on a Hypersil Gold C_18_, 1.9 μm, 2.1 mm × 100 mm column (Thermo Fisher) held at 40°C. Reversed phase liquid chromatography (RPLC) was performed at 250 μL min^-1^ using solvent A (0.1% FA in water) and solvent B (0.1% FA in acetonitrile/water 80:20) with a gradient length of 30 min as previously described^154, 155^. Intact MS analyses were acquired over the mass-to-charge ratio (m/z) range of 70-1,050 in positive ion mode at 120,000 resolution with a scan rate of 3.5 Hz, automatic gain control (AGC) target of 1x10^6^, and maximum ion injection time of 100 ms, and MS/MS (or tandem) spectra were collected at 15,000 resolution, AGC target of 2x10^5^ ions, with a maximum ion injection time of 100 ms.

### Metabolomics data processing and statistical analysis

Raw mass spectrometry (MS) data were imported, processed, normalized, and evaluated using Progenesis QI software (version 3.0; Non-linear Dynamics, Newcastle, UK). Relative abundances of spiked, heavy labeled QA/QC standards showed < 13% variability for sample preparation and <3% for instrument variability. All MS and MS/MS runs were aligned to a pooled quality control (QC) reference sample. Detected ions (i.e., unique retention time and m/z pairs), were de-adducted and de-isotoped to produce distinct analytical features. Data was normalized to all compounds, and statistical significance was evaluated using analysis of variance (ANOVA) on the normalized compound abundance values. Both tentative and putative metabolite annotations (Confidence Level 1-3)^156^ were assigned based on accurate mass measurements (mass error < 5 ppm), isotopic pattern similarity, MS/MS fragmentation spectrum, and retention time matching by querying the Human Metabolome Database^157^ and the CIT’s in-house reference library.

Metaboanalyst 5.0 (www.metaboanalyst.ca/) was used to perform high level visualizations including heat maps for compounds with statistical significance (p-value ≤ 0.05)^158^. The untargeted metabolomics data is available at the NIH Common Fund’s National Metabolomics Data Repository (NMDR) website, the Metabolomics Workbench, https://www.metabolomicsworkbench.org, under the assigned study where it has been assigned Project ID ST003919. The data can be accessed directly via its Project DOI: http://dx.doi.org/10.21228/M8PR93.

### 16S rDNA amplicon sequencing and analysis

Libraries were prepared from DNA isolated from cecal contents using Qiagen PowerSoil Pro Kit and the V3-V4 hypervariable region of 16S rRNA coding sequences was amplified. The library was constructed and sequenced using an Illumina MiSeq system, generating 250 bp paired-end, chimera-removed reads^159^. The downstream analysis was performed and visualized with QIIME2^160^ version 2024.5. Taxonomic profiling was performed against SILVA database version 132^161^. The sequencing reads generated during the current study are available at the European Nucleotide Archive repository under accession No. ENA: PRJEB89357 (secondary accession ENA: ERP172384).

### Bulk RNAseq and analysis

For *in vitro* bacterial RNAseq, ETBF was cultured in BHIS before subculture in lactate-supplemented BHIS anaerobically or in the presence of 0.1% O_2_ for 3 hours at 37 °C. Bacteria RNAprotect (Qiagen) was added to perverse the transcript. Bacteria were lysed using Lysing Matrix B (MP Bio) and Tri-reagent (Molecular Biology Center), and RNA was purified using the RNeasy mini kit (Qiagen). Paired-end, 150bp library was constructed using an RNAtag-seq strategy^142^ and sequenced on an Illumina NovaSeq (Illumina, CA). Reads were trimmed using BBMap software suite mapped to the ETBF genome (GCF_023702735.1_ASM2370273v1) using Bowtie2^142^. Mapped reads were quantified using featureCounts software package^162^ and differential expression analysis was performed using DESeq2 software^147^. The sequencing reads generated for this analysis are available at the European Nucleotide Archive repository under accession No. ENA: PRJEB89361 (secondary accession ENA: ERP172389).

For mouse tissue RNAseq, RNA was extracted from cecal tissues of ETBF infected mice using Tri-reagent (Molecular Biology Center) and RNeasy mini kit (Qiagen). Tissues were homogenized using Lysing Matrix C (MP Bio) and Tri-reagent (Molecular Biology Center), and RNA was purified using the RNeasy mini kit (Qiagen). The transcriptome was sequenced using the methods described above. Reads were mapped to the mouse genome (mm10, UCSC Genome Browser) using Bowtie2^142^, and differential expression analysis was performed using DESeq2 software^147^. Feature-level clustering, gene set enrichment analysis, and Weighted Gene Co-expression Network Analysis were performed on BigOmics Analytics ^163^. The sequencing reads generated for this analysis are available at the European Nucleotide Archive repository under accession No. ENA: PRJEB89362 (secondary accession ENA: ERP172390).

### Genome-Scale Metabolic Modeling

A genome-scale metabolic model for the enterotoxic strain of *B. fragilis* was reconstructed following established protocols^164^ and the COBRA toolbox version 2.13.3 for Matlab^165^. In particular, an existing manually-curated and data-validated GEM for *B. fragilis* strain 638R was used as a template^166^, along with several other gram-negative template GEMS from the BiGG database^167^: multiple *E. coli* GEMs^168, 169^, *Shigella boydii*^169^, *Shigella dysenteriae*^169^, *Shigella flexneri*^169^, and *Klebsiella pneumoniae*^170^. Metabolic genes used in each of these prior reconstructions were compared to genes in this strain of *B. fragilis* using BLAST (E-value < 10^-30^, identity > 50%); any matching genes and their associated reactions were added to a draft model then manually curated for accuracy. The model was curated to grow in defined media known to be suitable to *B. fragilis*, realistic ATP production, and mass balance as was done for the reconstruction of the existing 638R model^166^.

The model was then transformed into context-specific models for the analysis of fluxes in each of the three sample sites (cecum tissue, lumen, and mucus) utilizing StanDep^171^, an algorithm for deciding which reactions should be active in different conditions based on transcriptomic data. To simulate the complex medium of the human gut, all nutrients were provided to the models in low amounts (up to one millimole per gram dry weight per hour). The reduced models for each site were then optimized for maximal growth using parsimonious flux balance analysis^172^. This was then repeated for the isogenic ETBF mutant lacking *bft*. The underlying model for the mutant was not altered before StanDep processing as *bft* is not a metabolic gene present in the reconstruction.

### Quantification and statistical analysis

Unless noted otherwise, data analysis was performed in GraphPad Prism v10.4.2. Values of bacterial population sizes, competitive indices, fold changes in mRNA levels, and normalized colon length were normally distributed after transformation by the natural logarithm. A two-tailed Student’s *t*-test was used for ln-transformed data. Unless otherwise stated, ^∗^, *P* < 0.05; ^∗∗^, *P* < 0.01; ^∗∗∗^, *P* < 0.001; ns, not statistically significant. In all mouse experiments, *N* refers to the number of animals from which samples were taken. The outlined statistical details of experiments, including exact value of *N*, definition of center, and dispersion can be found in the figures and figure legends. Shapiro-Wilk test was used to determine the normality of log-transformed data. Sample sizes (i.e. the number of animals per group) were not estimated *a priori* since effect sizes in our system cannot be predicted. No predicted statistical outliers were removed since the presence or absence of these potential statistical outliers did not affect the overall interpretation. Mice that were euthanized early due to health concerns were excluded from analysis.

## Supporting information

Key Resource Table

## Acknowledgments

We thank Dr. James Imlay (University of Illinois Urbana-Champaign), Drs. Genevieve Mullins and Justin Milner (University of North Carolina at Chapel Hill) for insightful discussions and comments on the concepts of the work. R. T. F was supported by NIH (F31AI178950). Work in W. Z.’s lab was funded by the NIH (1R35GM147470 and 1R01DK134692), V Foundation (V2022-032), Colorectal Cancer Alliance (10065978), and The G. Harold & Leila Y. Mathers Charitable Foundation (MF-2207-03128). Work in M. G. B’s lab was funded by NIH (R35GM150625). Untargeted metabolomics work was supported in part using the resources of the Center for Innovative Technology (CIT) at Vanderbilt University. This work is supported by Metabolomics Workbench/National Metabolomics Data Repository (NMDR) (grant# U2C-DK119886), Common Fund Data Ecosystem (CFDE) (grant# 3OT2OD030544) and Metabolomics Consortium Coordinating Center (M3C) (grant# 1U2C-DK119889). Any opinions, findings, conclusions, or recommendations expressed in this material are those of the author(s) and do not necessarily reflect the views of the funding agencies. The funders had no role in study design, data collection, and interpretation, or the decision to submit the work for publication.

## Author contributions

L. S., R. T. F., A. Grote., A. Gnirke, A. M. E., and W. Z. designed the study. L. S. and R. T. F. performed and analyzed all the *in vitro* and *in vivo* experiments with help from M. L-B., A. K. M., D. S., A. E. R. and M. M. B. L. S., W. Z., A. Gnirke, B. B., and J. L. performed hybrid selection RNAseq and A. Grote performed all hybrid selection RNAseq analysis. N. M. and K. Z. performed genome-scale metabolic flux integration. M. W. C. performed targeted quantification of intestinal metabolites. A. C. S-R. and S. D. S. performed untargeted metabolomics and analyzed the data. O. F. H. and M. G. B. performed phylogenetic analysis of ETBF metabolic genes. L.S. performed florescence microscope imaging with the help from B. P. B.. M. K. W. quantified histopathological changes. L. S., D. O-V. and W. Z. performed SCENITH assay and immune cell proofing. L. S. analyzed *in vitro* RNAseq and 16S rDNA sequencing data with help from W. Z. L.S., R. T. F., A. Grote., A. M. E. and W. Z. interpreted the data, W. Z. wrote the manuscript, and all authors commented on the manuscript. For the untargeted metabolomic work, Investigation: S.G.C, S.D.S; Formal Analysis: A.C.S, S.D.S; Resources: S.D.S, J.A.M.; Project administration: S.D.S; Visualization: A.C.S; Writing – review and editing: S.G.C, A.C.S, S.D.S, J.A.M.

## Declaration of interests

The authors declared no financial interests.

---

**Fig. S1.**
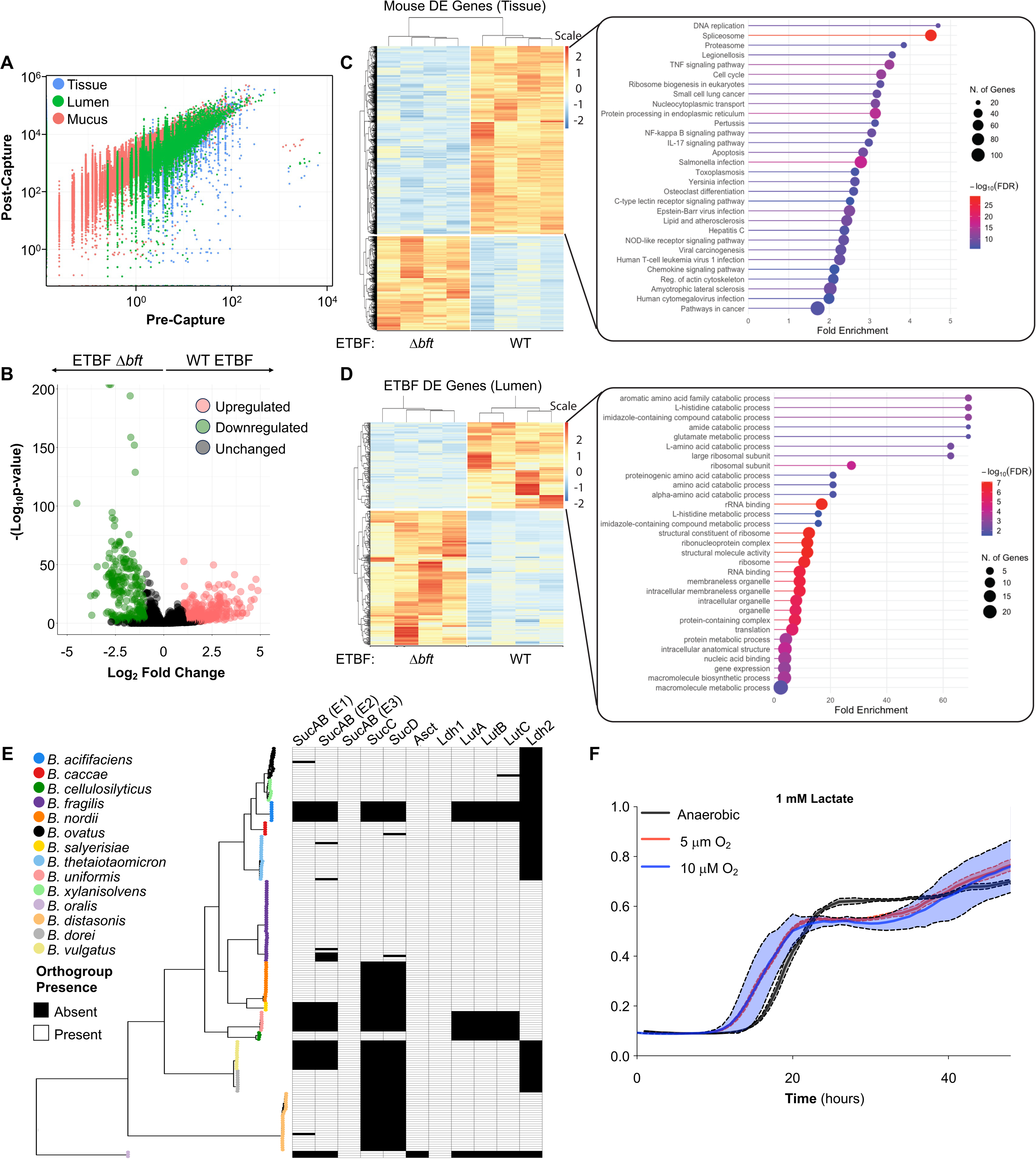
ETBF engages oxidative metabolism during intestinal colonization (related to Fig. 1). (**A-D**) C57BL/6 mice were colonized with either wild-type ETBF or an isogenic Δ*bft* mutant for 7 days. Hybrid-selection RNA-seq was performed on luminal contents, mucus, and cecal tissue to profile the ETBF transcriptome during infection. (**A**) Correlation between pre-captured and post-captured ETBF transcriptomes. (**B**) Volcano plot of post-captured ETBF transcriptome from the intestinal lumen. (**C-D**) Gene ontology (GO) enrichment analysis of (**C**) pre-captured mouse transcriptome and (**D**) post-captured ETBF transcriptome in the intestinal lumen. (**E**) Conservation of genes involved in lactate oxidation and central metabolism across the Bacteroidetes phylum. (**F**) The ETBF wild-type strain in semi-defined medium (SDM) supplemented with 1 mM lactate was exposed to indicated levels of O_2_. Bacterial growth was measured by optical density at 600 nm (OD_600_). Connected lines and shaded areas represent the mean and SEM.

**Fig. S2.**
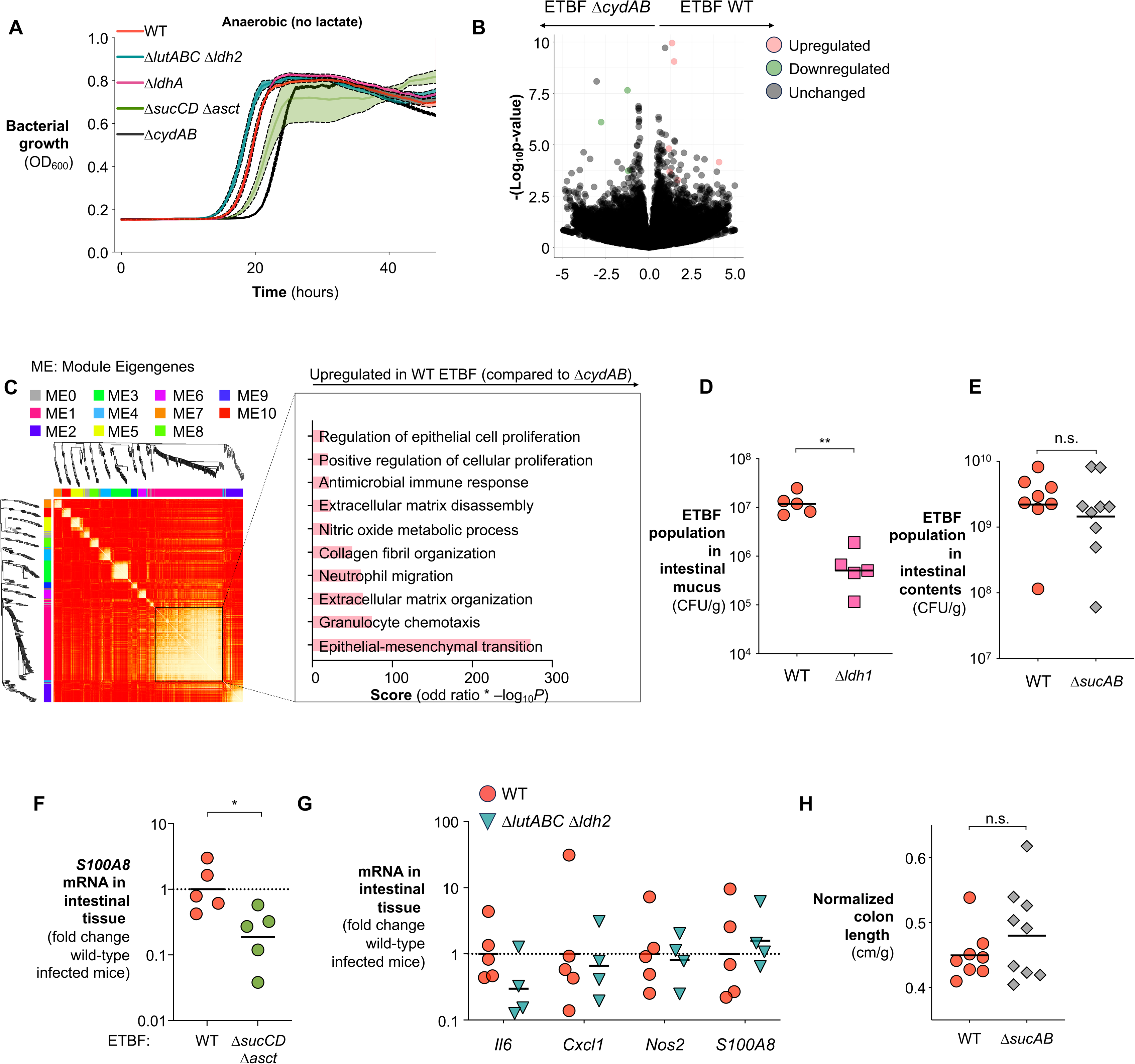
Oxygen respiration and oxidative metabolism contribute to ETBF colonization and inflammation (related to Fig. 2). (**A**) Indicated ETBF strains were cultured anaerobically in brain heart infusion (BHI) media for 48 hours, and bacterial growth was measured by optical density at 600 nm (OD_600_). Connected lines and shaded areas represent the mean and SEM. (**B-C**) C57BL/6 mice were colonized with either wild-type ETBF or the isogenic Δ*cydAB* mutant for 7 days. (**B**) Volcano plot of differentially expressed genes in the cecal transcriptome. (**C**) Weighted gene co-expression network analysis (WGCNA) of the cecal transcriptome. (**D-G**) C57BL/6 mice were colonized with either wild-type ETBF or the indicated mutant strains for 3 days. (**D**) Bacterial population in the intestinal mucus layer was quantified by plating on selective media. Transcript levels of inflammatory cytokines in intestinal tissue of mice colonized by (**E**) Δ*sucAB*, (**F**) Δ*sucCD* Δ*asct,* and (**G**) Δ*lutABC* Δ*ldh2* mutants were measured by RT-qPCR. Bars represent geometric means. n.s., not significant; *, *p*<0.05; **, *p*<0.01.

**Fig. S3.**
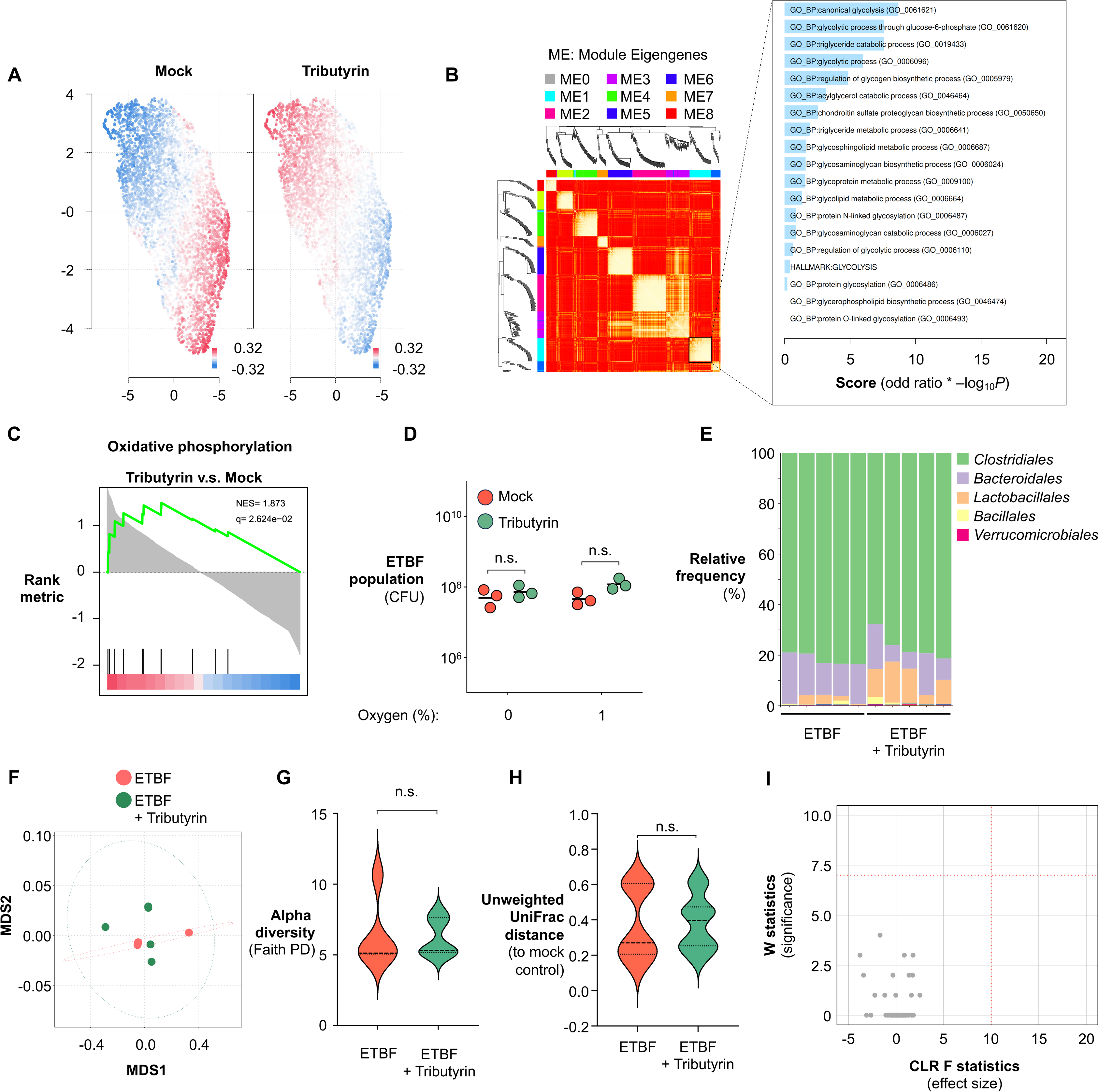
BFT-driven metabolic reprogramming of colonocytes promotes ETBF colonization (related to Fig. 3). (**A-C**) C57BL/6 mice colonized with ETBF were treated with either tributyrin or vehicle control for 7 days. (**A**) Feature-level clustering, (**B**) Weighted gene co-expression network analysis (WGCNA), and (**C**) Gene set enrichment analysis of the cecal transcriptome. (**D**) ETBF was cultured in brain heart infusion (BHI) medium supplemented with tributyrin and exposed to the indicated oxygen levels. Bacterial fitness was assessed by plating on selective agar. (**E-I**) C57BL/6 mice colonized with ETBF were treated with tributyrin or vehicle control. After 7 days, cecal contents were collected for DNA extraction and 16S rDNA sequencing to profile gut microbiota composition. (**E**) Order-level microbiota composition. (**F**) Multidimensional scaling (MDS) analysis of microbiota composition. (**G-H**) Violin plots showing (**G**) alpha diversity (Faith’s phylogenetic diversity) and (**H**) beta diversity (unweighted UniFrac distance). (**I**) Volcano plot from Analysis of Composition of Microbiomes (ANCOM), with dotted lines indicating minimal cutoff values for differentially abundant taxa. Bars represent geometric means. n.s., not significant. For **G**&**H**, the central thick dotted line indicates the median, while the upper and lower dotted lines correspond to the first and third quartiles (the 25th and 75th percentiles) of the data.

**Fig. S4.**
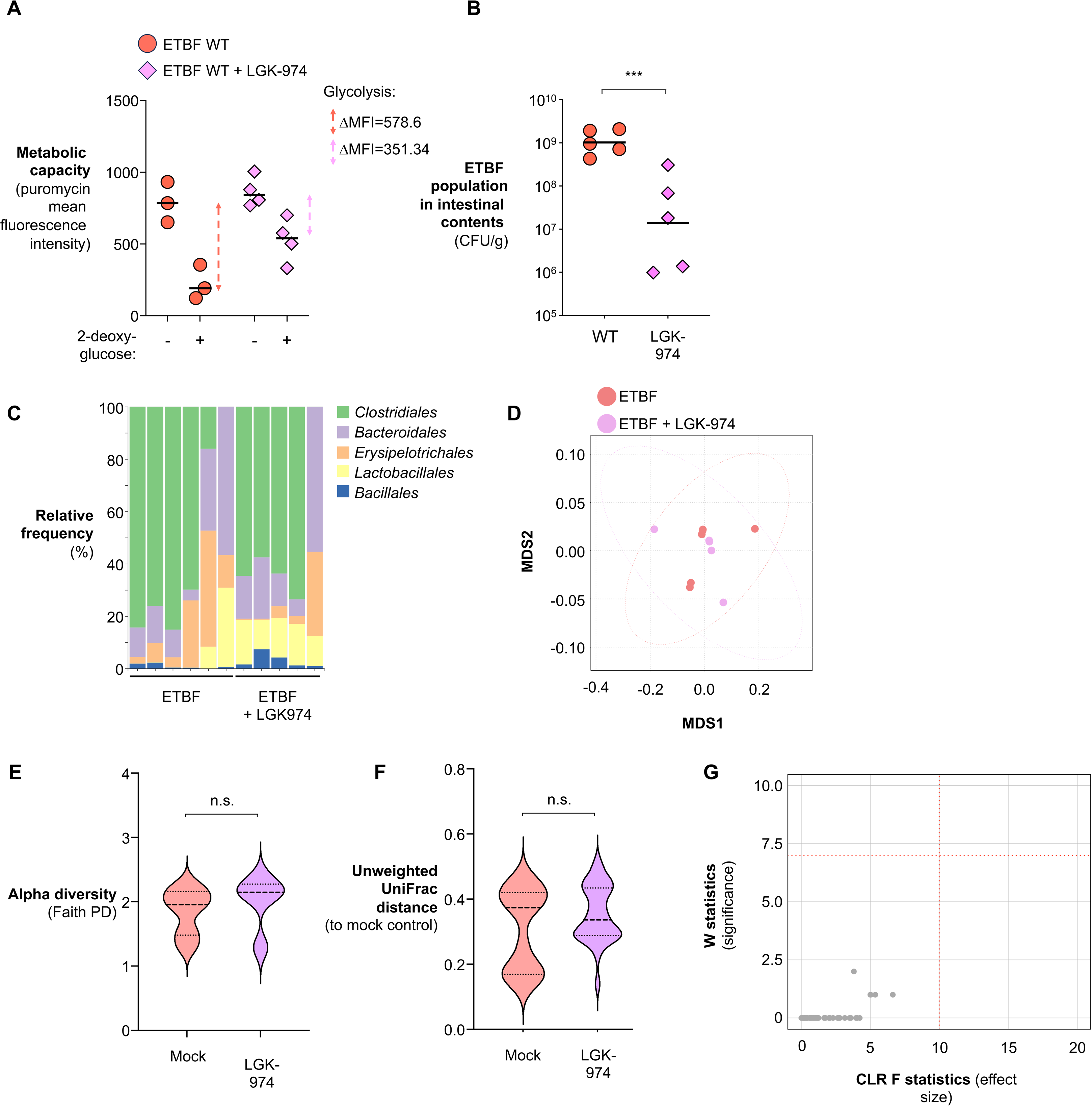
Disruption of epithelial processes promotes ETBF colonization and inflammation (related to Fig. 4). (**A**) C57BL/6 mice colonized with ETBF wild-type (WT) strain were treated with the Wnt/β-catenin inhibitor LGK-974 or vehicle control. After 7 days, metabolic activity and glycolytic dependency in gut epithelial cells was quantified using SCENITH assay. (**B)** C57BL/6 mice colonized with ETBF wild-type (WT) strain were treated with the Wnt/β-catenin inhibitor LGK-974 or vehicle control for 3 days. ETBF abundance was determined by plating on selective media. (**C-G**) C57BL/6 mice colonized with ETBF wild-type (WT) strain were treated with the Wnt/β-catenin inhibitor LGK-974 or vehicle control. After 7 days, cecal contents were collected, DNA was extracted, and 16S rDNA sequencing was performed to profile gut microbiota composition. (**C**) Order-level microbiota composition. (**D**) Multidimensional scaling (MDS) analysis of microbiota composition. (**E-F**) Violin plots showing (**E**) alpha diversity (Faith’s phylogenetic diversity) and (**F**) beta diversity (unweighted UniFrac distance). (**G**) Volcano plot from Analysis of Composition of Microbiomes (ANCOM), with dotted lines indicating minimal cutoff values for differentially abundant taxa. For **A**&**B**, bars represent geometric means. ***, *p*<0.001. For **E**&**F**, the central thick dotted line indicates the median, while the upper and lower dotted lines correspond to the first and third quartiles (the 25th and 75th percentiles) of the data.

**Fig. S5.**
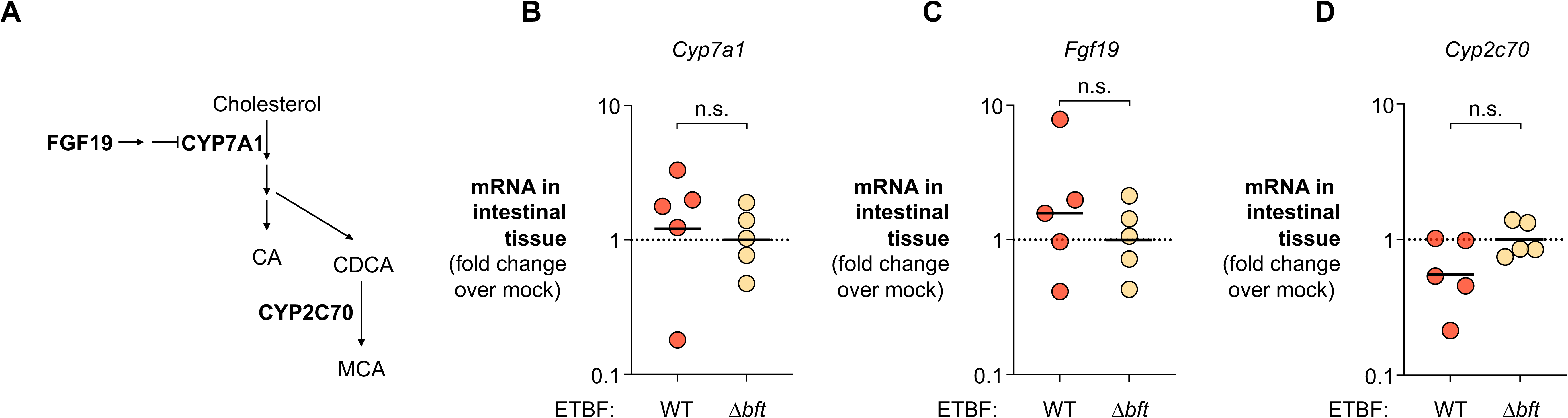
BFT depletes bile acids in the gut (related to Fig. 5). (**A-D**) C57BL/6 mice were colonized with either wild-type ETBF or the isogenic Δ*bft* mutant. 7 days later, liver tissue was collected and RNA extracted. (**A**) Schematic for regulation of bile acid biosynthesis pathway in the liver. (**B**) *Cyp7a1,* (**B**) *Fgf19*, and (**C**), *Cyp2c70* mRNA levels in the liver quantified by RT-qPCR. Bars represent geometric means. n.s., not significant.

**Fig. S6.**
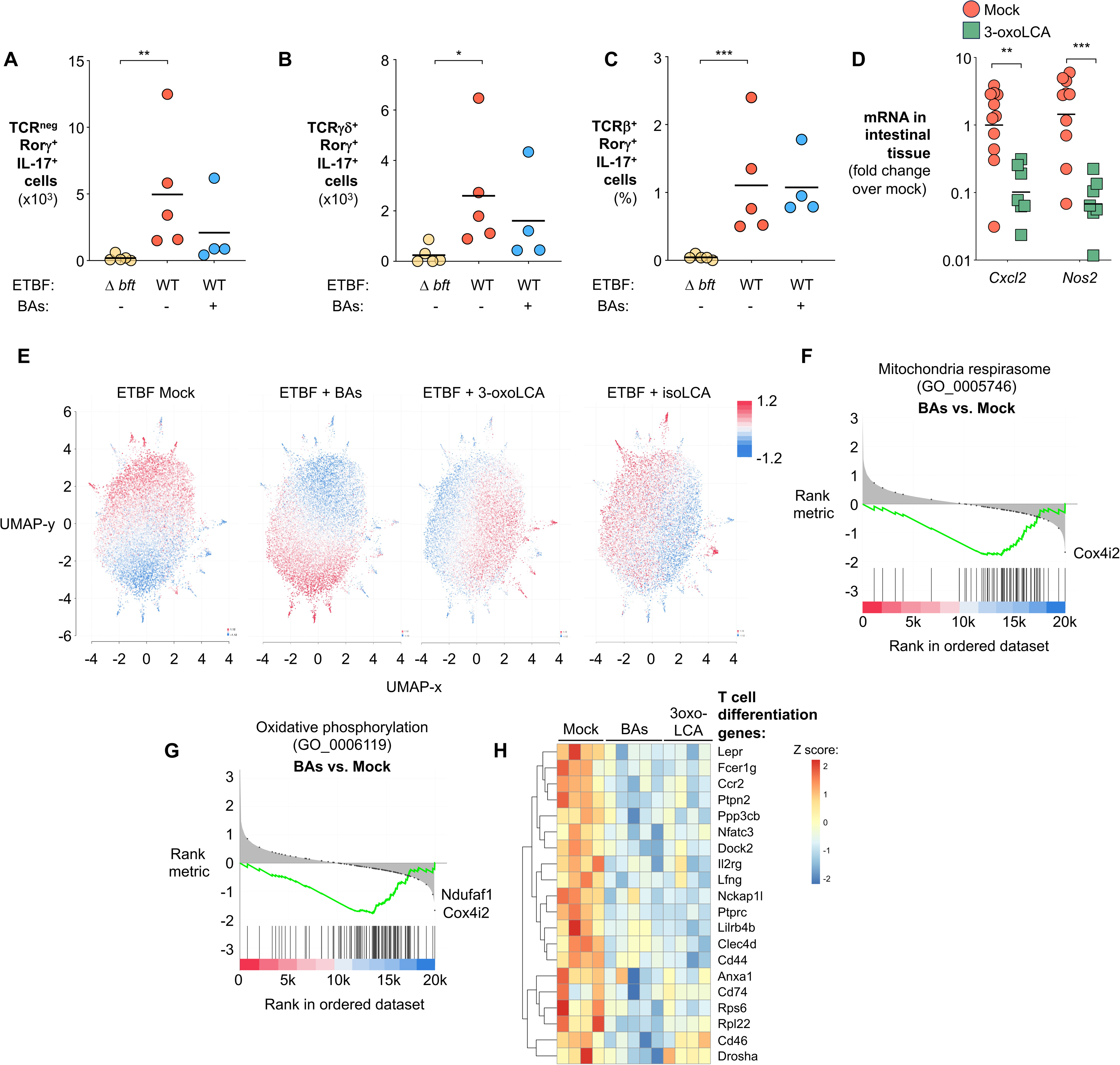
ETBF rewires colonocyte metabolism by impacting bile acid abundance (related to Fig. 6). (**A-C**) Groups of C57BL/6 mice were colonized with wild-type ETBF or the isogenic Δ*bft* mutant and given either vehicle control or primary bile salts in drinking water for 7 days. IL-17-expressing intraepithelial lymphocytes were profiled by flow cytometry. Quantification of IL-17-producing (**A**) innate lymphoid cells and (**B**) γδ T cells, and the frequency of (**C**) TH17 cells. (**D-H**) Groups of ETBF-colonized C57BL/6 mice were administered the indicated bile acids intragastrically for 7 days. (**D**) mRNA levels of inflammatory cytokines in intestinal tissue determined by RT-qPCR. (**E**) feature-level clustering, (**F&G**) gene set enrichment analysis, and (**H**) Pathway analysis of the cecal transcriptome. Bars represent geometric means. *, *p*<0.05; **, *p*<0.01; ***, *p*<0.001.

**Fig. S7.**
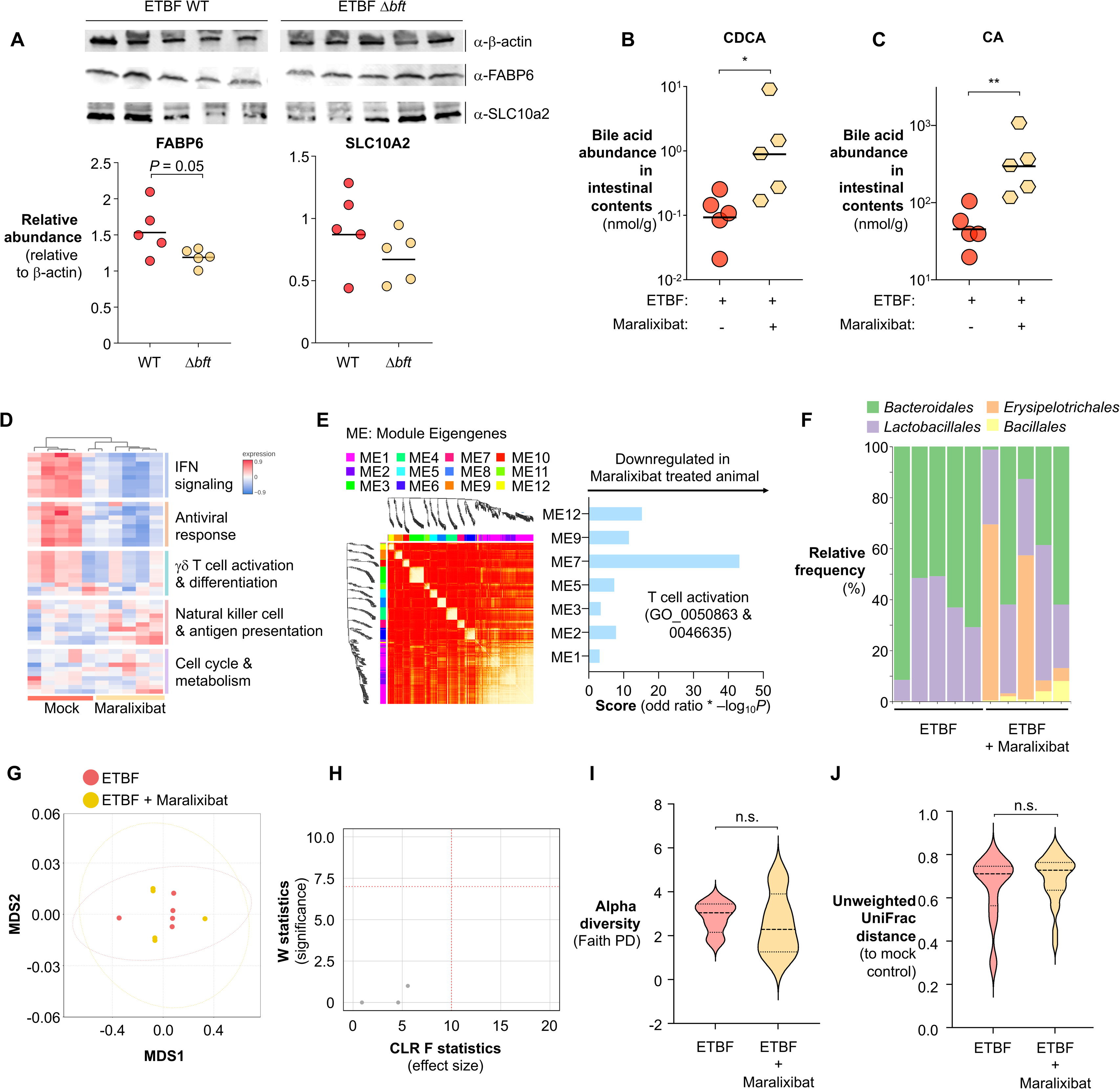
ETBF exploits the bile acid recycling pathway to promote bile acid depletion and gut inflammation (related to Fig. 7). **(A)** C57BL/6 mice were colonized with either wild-type ETBF or the isogenic Δ*bft* mutant for 7 days. SLC10A2 and FABP6 protein levels in the colonic tissue were assessed using specific antibodies. Relative abundance was quantified in the panel below. **(B-J)** C57BL/6 mice colonized with ETBF were treated with either a Maralixibat-fortified diet or a control diet for 7 days. **(B-C**) Bile acid levels in intestinal contents measured by LC-MS/MS: **(B)** chenodeoxycholic acid (CDCA) and (C) cholic acid (CA). **(D-E)** Intestinal RNA was extracted and transcriptome profiled by RNA-seq. (D) Heatmap of differentially expressed genes and enriched signaling pathways. **(E)** Weighted gene co-expression network analysis (WGCNA) of the cecal transcriptome. **(F-J)** Gut microbiota composition was analyzed by 16S rDNA sequencing of intestinal contents. (F) Order-level microbiota composition. **(G)** Multidimensional scaling (MDS) analysis of microbiota composition. **(H)** Volcano plot from Analysis of Composition of Microbiomes (ANCOM), with dotted lines indicating minimal cutoffs for differential abundance. **(I)** Violin plot showing alpha diversity (Faith’s phylogenetic diversity). **(J)** Violin plot showing beta diversity (unweighted UniFrac distance). Bars represent geometric means. n.s., not significant; *, *p*<0.05; **, *p*<0.01. For I&J, the central thick dotted line indicates the median, while the upper and lower dotted lines correspond to the first and third quartiles (the 25th and 75th percentiles) of the data.

**Fig. S8.**
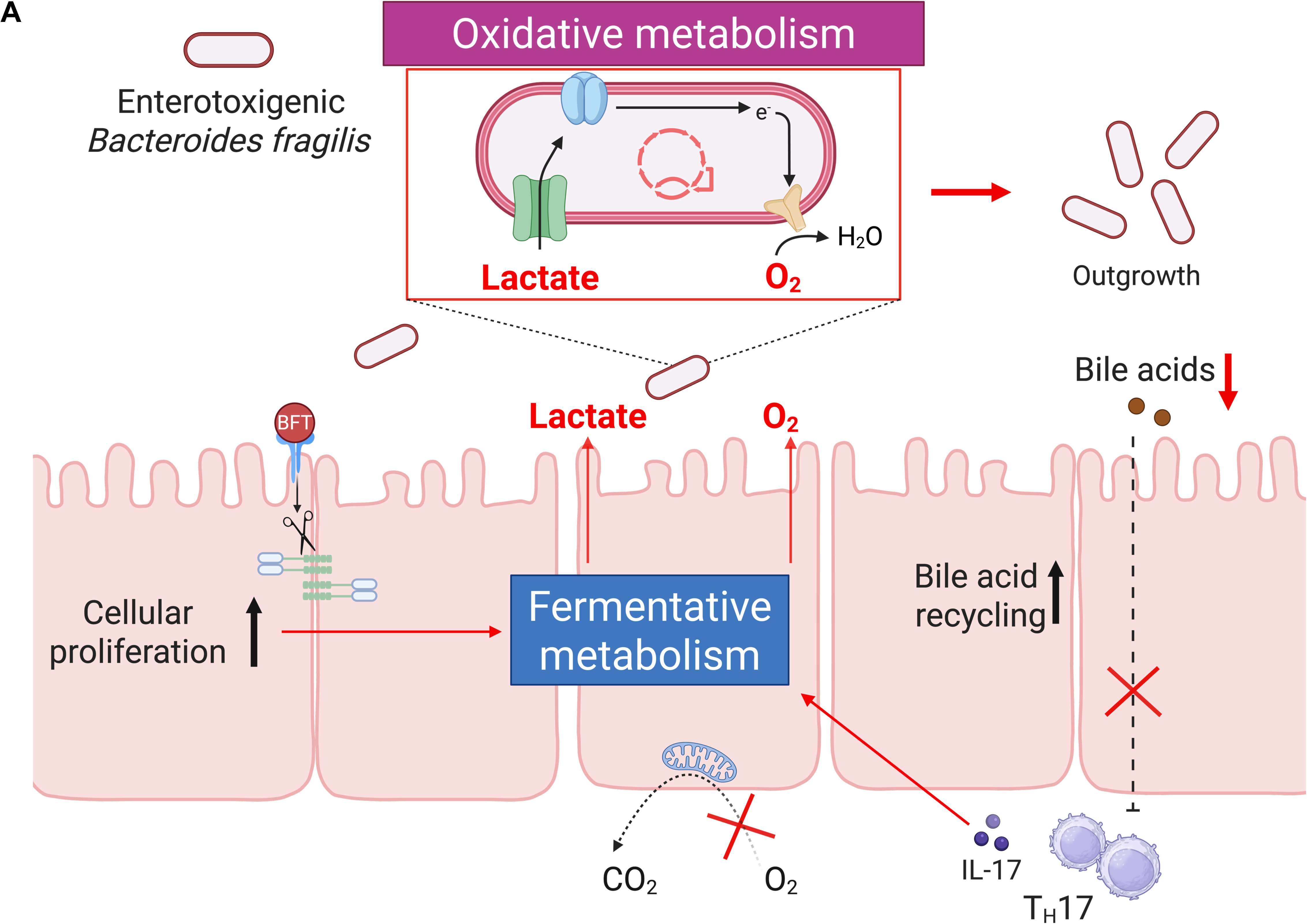
Proposed model for ETBF colonization of the inflamed gut (related to Figs. 1-7). (**A**) We propose that BFT induces a metabolic shift in colonocytes from oxidative phosphorylation to fermentative metabolism, resulting in increased O_2_ and lactate levels in the gut. These alterations create an oxidative environment within the otherwise anaerobic gut, enabling ETBF, an obligate anaerobe, to operate an oxidative metabolism to colonize the inflamed gut.

## Notes

### Competing Interest Statement

The authors have declared no competing interest.

