## Supplementary material for "An Anaerobic Pathogen Rewires Host Metabolism to Fuel Oxidative Growth in the Inflamed Gut": Key Resource Table

**KEY RESOURCES TABLE**

| REAGENT or RESOURCE | SOURCE | IDENTIFIER AND USAGE |
| --- | --- | --- |
| **Bacterial Strains and Quantification Method** | | |
| *Enterotoxigenic Bacteroides fragilis* 86-5443-2-2 | [1] | WZ1023 |
| *B. fragilis* Δ*bft* | [2] | WZ1025 |
| *B. fragilis* Δ*tdk* | This study | RF54 |
| *B. fragilis* Δ*tdk* Δ*bft* | This study | RF59 |
| Non-enterotoxigenic *B. fragilis* ATCC 25285 |  | WZ901 |
| *B. fragilis* Δ*tdk* Δ*cydAB* | This study | RF252 |
| *B. fragilis* Δ*tdk* Δ*lutABC* Δ*ldh2* | This study | LS82 |
| *B. fragilis* Δ*tdk* Δ*ldh1* | This study | RF314 |
| *B. fragilis* Δ*tdk* Δ*sucCD* Δ*asct* | This study | WZ1481 |
| *B. fragilis* Δ*tdk* Δ*sucAB* | This study | LS80 |
| E. coli S17-1 λpir; zxx::RP4 2-(Tet^r^::Mu) (Kan^r^::Tn7) λpir |  | S17-1 λpir |
| **Recombinant DNA** | | |
| pKNOCK-*bla*-*ermGb* | [3] | pKNOCK |
| pKNOCK-*bla*-*ermGb*::*tdk* | [3] | pExchange-tdk |
| pKNOCK-*bla*::Promoter *ermGb*::*cfxA* | This study | pKNOCK-ETBF |
| pKNOCK-*bla*::Promoter *ermGb*::*cfxA*::*tdk* | This study | pEET |
| pNBU2-*bla*-*catP* | This study | pNBU2-CmR |
| Upstream and downstream regions of ETBF *cydAB* in pEET | This study | pRF237 |
| Upstream and downstream regions of ETBF *lutABC* in pEET | This study | pRF231 |
| Upstream and downstream regions of ETBF *asct* in pEET | This study | pML191 |
| Upstream and downstream regions of ETBF *ldh2* in pEET | This study | pML108 |
| Upstream and downstream regions of ETBF *ldh1* in pEET | This study | pML102 |
| Upstream and downstream regions of ETBF *sucAB* in pEET | This study | pRF317 |
| Upstream and downstream regions of ETBF *sucCD* in pEET | This study | pRF247 |
| **Oligonucleotides** | | |
| Δ86_*tdk* 1 up:  5’- gctctagaactagtggatccGCTTTACAAGAAGATCGAG-3’ | This study | N/A |
| Δ86_*tdk* 1 down:  5’-gagctcccaTTGTATATGATCTTCTGAAAATAATAC-3’ | This study | N/A |
| Δ86_*tdk* 2 up:  5’-tgccatttatTTGTATATGATCTTCTGAAAATAATAC-3’ | This study | N/A |
| Δ86_*tdk* 2 down:  5’-tcgaattcctgcagcccgggATTCCTGAGTCAAAGACTC-3’ | This study | N/A |
| ETBF Δtdk verification primer F:  5’- AGCGAGGATTTAAGACTGTA-3’ | This study | N/A |
| ETBF Δtdk verification primer R:  5’- GTTCCTCAATACGTTGAGTT-3’ | This study | N/A |
| Δ86_*lutABC* 1 up:  5’-gctctagaactagtggatccTGCACATAAGCTGATTCTTC-3’ | This study | N/A |
| Δ86_*lutABC* 1 down:  5’-gagctcccatATAAATGGCATTGATATAACAGG-3’ | This study | N/A |
| Δ86_*lutABC* 2 up:  5’-tgccatttatATGGGAGCTCATGGAGCG-3’ | This study | N/A |
| Δ86_*lutABC* 2 down:  5’-tcgaattcctgcagcccgggCAGAGTCACTACTTAAATCAACTCGTAG-3’ | This study | N/A |
| Δ86_*lutABC* verification primer F:  5’-CCAACGGGATCAGGAAGAAA-3’ | This study | N/A |
| Δ86_*lutABC* verification primer R:  5’-ACAGACGTGACTGTGGTATTG-3’ | This study | N/A |
| Δ86_*ldh2* 1 up:  5’- ggccgctctagaactagtggATAAATAAGGGGTTGACACC-3’ | This study | N/A |
| Δ86_*ldh2* 1 down:  5’- agatatcggaTGTACCCGGCCTATAAAAC-3’ | This study | N/A |
| Δ86_*ldh2* 2 up:  5’- gccgggtacaTCCGATATCTTGCAAGAAC-3’ | This study | N/A |
| Δ86_*ldh2* 2 down:  5’- gaattcctgcagcccgggGGATTCACTTTACGGAAAGAC-3’ | This study | N/A |
| Δ86_*ldh2* verification primer F:  5’- TGCGAATGCCAGAGAACCGG-3’ | This study | N/A |
| Δ86_*ldh2* verification primer R:  5’- TTAATATCCACCATTGACAT-3’ | This study | N/A |
| Δ86_*ldh1* 1 up:  5’- gctctagaactagtggatccCTCGAAATGTTTTGATTTGAAAATC-3’ | This study | N/A |
| Δ86_*ldh1* 1 down:  5’- ttatgatgaaAATCAGAAGCCTTTGGTG-3’ | This study | N/A |
| Δ86_*ldh1* 2 up:  5’- gcttctgattTTCATCATAAGGTTTGGTAC-3’ | This study | N/A |
| Δ86_*ldh1* 2 down:  5’- tcgaattcctgcagcccgggTTGGCATAATCACTGAAATAC-3’ | This study | N/A |
| Δ86_*ldh1* verification primer F:  5’- GAAAAAATGAGAGAATAACC-3’ | This study | N/A |
| Δ86_*ldh1* verification primer R:  5’- GATAAGCATCTGCGGCTGTTC-3’ | This study | N/A |
| Δ86_*sucCD* 1 up:  5’- tagtggatccTTCAACGGTACGGACGAAG-3’ | This study | N/A |
| Δ86_*sucCD* 1 down:  5’- agggttcaccGTAAGTGGAGAAAATCTCCTTTG-3’ | This study | N/A |
| Δ86_*sucCD* 2 up:  5’- ctccacttacGGTGAACCCTCGGAAATAC-3’ | This study | N/A |
| Δ86_*sucCD* 2 down:  5’- gcagcccgggCGAAGCAAACCAATGAAATG-3’ | This study | N/A |
| Δ86_*sucCD* verification primer F:  5’- GTATTCCCCGGTCAAGGTGC-3’ | This study | N/A |
| Δ86_*sucCD* verification primer R:  5’- CTCCAATTGTCTCTAATTCC-3’ | This study | N/A |
| Δ86_*asct* 1 up:  5’-cgctctagaactagtggatccATGGTGAAAGTGACAGC-3’ | This study | N/A |
| Δ86_*asct* 1 down:  5’-cagccttgtcAGCGGAGACATGAAAAAC-3’ | This study | N/A |
| Δ86_*asct* 2 up:  5’-tgtctccgctgacaaggCTGGCAGCTTC-3’ | This study | N/A |
| Δ86_*asct* 2 down:  5’-cgaattcctgcagcccgggGTTTTCTTCTTCATCGAATGAAAGAACTC-3’ | This study | N/A |
| Δ86_*asct* verification primer F:  5’-TTACTTTCCCTTTCTCGGTATTG-3’ | This study | N/A |
| Δ86_*asct* verification primer R:  5’-TCACCGGCCTCTACCGAAGC-3’ | This study | N/A |
| Δ86_*sucAB* 1 up:  5’-cgctctagaactagtggatcTTGACCTGCACCCTTAACTCTAC-3’ | This study | N/A |
| Δ86_*sucAB* 1 down:  5’-tcttttcatcATACCCGGCAGGTCCACC-3’ | This study | N/A |
| Δ86_*sucAB* 2 up:  5’-tgccgggtatgatgaaAAGATATATAAAGCAGCCAAAGAGTTGC-3’ | This study | N/A |
| Δ86_*sucAB* 2 down:  5’-cgaattcctgcagcccgggGATGCGGGGAGCTAAGCCTTG-3’ | This study | N/A |
| Δ86_*sucAB* verification primer F:  5’- TCGTTTCTGTGGCTATCACA-3’ | This study | N/A |
| Δ86_*sucAB* verification primer R:  5’- CCTGATCAATCCGCAGATGG-3’ | This study | N/A |
| Mouse *Gapdh* qRT-PCR primers:  5’-TGTAGACCATGTAGTTGAGGTCA-3’  5’-AGGTCGGTGTGAACGGATTTG-3’ | [4] | N/A |
| Mouse *Cxcl1* qRT-PCR primers:  5’-TGCACCCAAACCGAAGTCAT-3’  5’-TTGTCAGAAGCCAGCGTTCAC-3’ | [5] | N/A |
| Mouse *IfnG* qRT-PCR primers:  5’- TCAAGTGGCATAGATGTGGAAGAA-3’  5’- TGGCTCTGCAGGATTTTCATG -3’ | [4] | N/A |
| Mouse *Il17* qRT-PCR primers:  5’- ATCGCTGCTGCCTTCACTGTAG-3’  5’- TGATGCTGTTGCTGCTGCTGAG-3’ | This study | N/A |
| Mouse *Fgf15* qRT-PCR primers:  5’- ATGGCGAGAAAGTGGAACGG -3’  5’- CTGACACAGACTGGGATTGCT -3’ | This study | N/A |
| Mouse *Cyp7a1* qRT-PCR primers:  5’- CACCTTGAGGATGGTTCCTATAAC -3’  5’- CCAAAGGGTCTGGGTAGATTTC -3’ | This study | N/A |
| Mouse *Cyp2c70* qRT-PCR primers:  5’- CAAGTCATCTCCTCCACCATAC -3’  5’- CCAGGTTTCAGCACCAAATAAA -3’ | This study | N/A |
| Mouse *Cyp2a12* qRT-PCR primers:  5’- CTGAGAACAAAGCCCTCTACTT -3’  5’- CAACTTTCAGGAGGACCATACA -3’ | This study | N/A |
| **Chemicals, Peptides, and Recombinant Proteins** |  |  |
| (NH_4_)_2_SO_4_ | Sigma | A4915 |
| 2-Deoxy-D-glucose ≥99% | Fisher Scientific | AC111980050 |
| 5 beta -CHOLANIC ACID-3-ONE | Steraloids | C1750-000-50mg |
| 5-fluoro-2-deoxy-uridine | Ark Pharm | AK-24802 |
| Acid-Phenol:Chloroform:IAA, pH 4.5 (125:24:1) | Ambion | AM9720 |
| Agar | Thermo Fisher | BP1423 |
| Anti rabbit IgG (H+L), F(ab')2 Fragment (Alexa Fluor® 488 Conjugate) | Cell Signaling Technology | 4412S |
| Anti-CD326 (Ep-CAM) Rat Monoclonal Antibody (APC (Allophycocyanin)) [clone: G8.8], Size=100 μg | BioLegend | 118214 |
| Anti-Mouse CD45, PE-Cyanine7 (30-F11) | TonBo Biosceinces | 60-0451-U025 |
| Anti-Mouse ROR ˠt (B2D) | Invitrogen | 12-6981-80 |
| Anti-Mouse TCR-beta, APC Cyanine7 (H57-597) | Cytek | 25-5961-U025 |
| Anti-SLC10A2 Rabbit Polyclonal Antibody, Size=150 µL | Proteintech | 25245-1-AP |
| Bile salts | Sigma-Aldrich | B8756-100G |
| BioReagent Puromycin dihydrochloride, ≥98% (HPLC), From Streptomyces alboniger | Sigma-Aldrich | P8833-100MG |
| Biotin Anti-Mouse TCR ˠ/𝛿 (GL3) | BioLegend | 118103 |
| Brain-heart-infusion (BHI) media | Becton Dickinson | 211059 |
| Carbenicillin disodium salt | VWR | J358-1G |
| Chenodeoxycholic acid | Cambridge Isotope Laboratories | ULM-9540-0.05 |
| Chenodeoxycholic acid (2,2,4,4-d4, 98%) | Cambridge Isotope Laboratories | DLM-6780-0.05 |
| Chloroform, anhydrous, >99% | Sigma-Aldrich | 288306-1L |
| Cholic acid | Cambridge Isotope Laboratories | ULM-9543-0.05 |
| Cholic acid (2,2,4,4-D4, 98%) | Cambridge Isotope Laboratories | DLM-2611-0.05 |
| Clindamycin Hydrochloride Monohydrate | TCI Chemicals | C2256-25G |
| Collagenase from *Clostridium histolyticum* | Sigma-Aldrich | C2139-500MG |
| Dansyl hydrazine | Sigma | 635928-500MG |
| Deoxycholic Acid (2,2,4,4-d4, 98%) | CDN Isotopes | D-2941 |
| Difco LB broth-Miller | Becton Dickinson | Cat# 244620 |
| 1-(3-Dimethylaminopropyl)-3-ethy;carbodiimide hydrochloride, >98% | Thermo | A10807.06 |
| Dimethyl sulfoxide (DMSO) | Corning | MT-25950CQC |
| DL-Dithiothreitol solution,1 m in H2O | Sigma-Aldrich | 646563-0X.5ML |
| DL-Dithiothreitol solution,1 m in H2O | Sigma-Aldrich | 646563-10X.5ML |
| DNA-free DNA removal Kit | Invitrogen | AM1906 |
| DNase 1 | Sigma | D4527-20KU |
| Dry Powder Milk, nonfat | RPI | M17200-500 |
| EDTA Disodium Salt | RPI | E57020-1000.0 |
| Erythromycin | Sigma-Aldrich | E5389-1G |
| FABP6 Polyclonal antibody | Proteintech | 13781-1-AP |
| Fetal Bovine Serum, 500 mL, Regular, USDA Safety Tested (Heat Inactivated) | Corning | 35-011-CV |
| Gentamicin | Sigma-Aldrich | G1397-100ML |
| Ghost Dye(tm) Violet 510 Viability Dye | Cell Signaling Technology | 59863S |
| Girard’s Reagent P | TCI Chemicals | G0030 |
| Glucose | Fisher Chemical | D14-212 |
| Glyceryl tributyrate,≥99% | Sigma-Aldrich | T8626-100ML |
| Glycocholic Acid (2,2,4,4-d4, 98%) | CDN Isotopes | D-3878 |
| Goat Anti-Mouse IgG Polyclonal Antibody (IRDye 800CW), Size=500 µg | Neta Scientific | 926-32210 |
| GolgiStop protein transporter inhibitor | BD Bioscience | 554724 |
| Hemin, from Porcine, >96% (HPLC) | Sigma | 51280-1G |
| HBSS (10X), no calcium, no magnesium, no phenol red | Thermo Scientific | 14185052 |
| HBSS, calcium, magnesium | Thermo Scientific | 24020117 |
| Hypoxyprobe Omni Kit (200 mg pimonidazole HCl plus 1 unit of 2627 rabbit antisera) | Hypoxyprobe | HP3-200Kit |
| Hypoxyprobe-Green Kit (100 mg pimonidazole HCl plus 1 unit of 4.3.11.3 mouse FITC-MAb) | Hypoxyprobe | HP6-100Kit |
| Isolithocholic Acid | Cayman Chemical Company | 29545 |
| Kanamycin | Fisher Chemical | BP906-5 |
| KH_2_PO_4_ | Fisher Chemical | P285 |
| L-methionine | Sigma | M9625 |
| LGK-974 5mg | Selleck Chemicals | S7143 |
| LI-COR IRDye 800cw Donkey Anti-Rabbit Igg Secondary Antibody, 0.1 mg | Neta Scientific | 925-32213 |
| Lithocholic acid | Cambridge Isotope Laboratories | ULM-9559-0.05 |
| Lithocholic acid (2,2,4,4-D4, 98%) | Cambridge Isotope Laboratories | DLM-9560-0.05 |
| Medchemexpress LLC MARIMASTAT 5MG SDP | Fisher Scientific | 50-187-2596 |
| Menadione (Vitamin K_3_) | Sigma | M5625-25G |
| Methanol, anhydrous, 99.8% | Sigma-Aldrich | 322415-1L |
| Mouse Cot-1 DNA, 1µg/µL | Thermo | 18440016 |
| NaCl | RPI | S23020 |
| NaHCO_3_ | Sigma | S6014 |
| Oligomycin A, ≥99% (HPLC) | Sigma-Aldrich | 75351-5MG |
| Paraformaldehyde Aqueous Solution, 16% | Electron Microscopy Sciences | 15710 |
| PE anti-Puromycin Antibody | BioLegend | 381504 |
| Percoll | Cytiva | 17089101 |
| Proteinase K, Molecular Biology Grade, 800 U/mL | New England Biolabs (NEB) | P8107S |
| Protoporphyrin IX | Sigma | P8293 |
| Pyridine Hydrochloride, 98% | Sigma-Aldrich | 243086 |
| Rat Anti-Mouse Foxp3, Alexa Fluor®️ 647 (MF23) | BD Pharmingen | 560401 |
| Rat Anti-Mouse IL-17A, BV421 (TC11-18H10) | BD Horizon | 563354 |
| redFluor™ 710 Anti-Mouse CD45 (30-F11) | Tonbo Biosciences | 80-0451-U100 |
| Resazurin | Sigma | R7017 |
| RIPA Lysis and Extraction Buffer | Thermo Scientific | 89900 |
| RPMI Medium 1640 (1X) | gibco | 22400-089 |
| Sodium Dodecyl Sulfate, 20% | Invitrogen | AM9820 |
| Sodium L-lactate,≥99.0% | Sigma | 71718-10G |
| Streptavidin PE-eFluor 610, eBioscience | Invitrogen | 61-4317-82 |
| Streptomycin Sulphate USP Grade | G Biosciences | RC-196 |
| Tcr gamma/delta Monoclonal Antibody (eBioGL3 (GL-3, GL3)), eFluor 450, eBioscience | Thermo Scientific | 48-5711-82 |
| TRI reagent | Molecular research center | TR118 |
| Tryptone | Thermo Fisher | BP1421 |
| Ursodeoxycholic Acid (2,2,4,4-d4) | CDN Isotopes | D-3819 |
| α-pimonidazole rabbit antisera | Hypoxyprobe | Pab2627 |
| **Critical Commercial Assays** |  |  |
| 384-well Low Flange White Flat Bottom Polystyrene TC-treated Microplates, with Lid, Sterile | Corning | 3570 |
| Bacteria RNAprotect | Qiagen | 76506 |
| BacTiter-Glo Microbial Cell Viability Assay | Promega | G8230 |
| Foxp3 / Transcription Factor Staining Buffer Set | Thermo Scientific | 00-5523-00 |
| Gibson Assembly Cloning Kit | NEB | E2611 |
| KAPA HiFi Hotstart ReadyMix | Roche | KK2601 |
| Lysing Matrix B | MP Bio | 1169110-CF |
| Lysing Matrix C | 1169120-CF | MP Bio |
| M.O.M. (Mouse on Mouse) Immunodetection Kit, Basic | VECTOR laboratories | BMK-2202 |
| Mini-Protean 4-20% TGX Stain-Free Gel | Bio-Rad | 4568095 |
| PowerSoil Pro Kit | Qiagen | 47014 |
| PowerUp SYBR Green Master Mix | Applied Biosystems | A25742 |
| Q5 Hot Start 2x Master Mix | NEB | M0494L |
| RNAlater™ Stabilization Solution | Thermo Fisher | AM7020 |
| RNeasy mini kit | Qiagen | 74104 |
| SuperScript VILO cDNA Synthesis Kit | Invitrogen | 11754050 |
| SYBR Green qPCR Master Mix | Life Technologies | 4309155 |
| TaqMan reverse transcription reagents | Invitrogen | N8080234 |
| Trans-Blot Turbo RTA Mini 0.2 µm PVDF Transfer Kit | Bio-Rad | 1704272 |
| Turbo DNA-free Kit | Ambion | AM1907 |
| Twist Standard Hybridization Reagents Kit, v1 | Twist Bioscience | 104179 |
| **Deposited Data** |  |  |
| 16S rRNA sequencing | The European Nucleotide Archive | ERP172384 |
| **Experimental Models: Organisms/Strains** |  |  |
| SPF C57BL/6 mice (wild-type) | The Jackson Laboratory | Cat# 000664 |
| **Software and Algorithms** |  |  |
| BBMap | DOE Joint Genome Institute | V38.90 |
| BigOmics Analytics | [6] | V3.2.26 |
| BioRender | BioRender.com | N/A |
| Bowtie2 | [7] | V2.5.4 |
| CFX Maestro | Bio-Rad | V2.3 |
| COBRA | MATLAB | V2.13.3 |
| DESeq2 | [8] | V1.48.0 |
| Excel for Mac | Microsoft | V16.70 |
| FlowJo | FlowJo | V10.10 |
| Ggtree | [9] | V3.16.0 |
| Image J | Fiji | 1.54p |
| MetaboAnalyst 5.0 | [10] | V5.0 |
| NCBI Datasets Command Line Tool | PMID: 38969627 | V16.40.1 |
| NIS-Elements | Nikon | V6.10.01 |
| Odyssey Western Blot Image Studio | LI-COR | V6.0 |
| OrthoFinder | [11] | V3.0.1b1 |
| Prism | Graph Pad | V10.4.2 |
| Progenesis QI | Non-Linear Dynamics | V3.0 |
| QIIME2 | [12] | V2024.5 |
| SILVA Database | [13] | V132 |
| Thermo Xcalibur | Thermo | V2.0.7 SP1 |
| Thermo LCQuan | Thermo | V2.7 |
